# Kinesin-1 is highly flexible and adopts an open conformation in the absence of cargo

**DOI:** 10.1101/2024.12.20.629623

**Authors:** Evelyn R. Smith, Emma D. Turner, Mahmoud A. S. Abdelhamid, Timothy D. Craggs, Alison E. Twelvetrees

**Affiliations:** Division of Neuroscience, School of Medicine and Population Health, Faculty of Health, The University of Sheffield, Sheffield Institute for Translational Neuroscience, 385a Glossop Road, Sheffield, S10 2HQ, UK; School of Mathematical and Physical Sciences, Faculty of Science, The University of Sheffield, Dainton Building, Brook Hill, S3 7HF, UK; Exciting Instruments, The Innovation Centre, 217 Portobello, Broomhall, Sheffield S1 4DP, UK

## Abstract

Kinesin-1 is an essential anterograde microtubule motor protein. The core kinesin motor is a homodimer of two heavy chains; N-terminal motor domains hydrolyse ATP and walk along microtubules, whilst a long elongated coiled-coil stalk and an intrinsically disordered C-terminal tail region bind cargos. Kinesin autoinhibition is key to preventing futile ATP consumption and occurs, at least in part, through direct interactions between N-terminal motor domains and C-terminal inhibitory motifs. Despite significant advances in our understanding of kinesin walking, little is known about the kinesin-1 conformational landscape of the stalk and tail domains. Here we apply solution based biophysical analysis tools to study conformational changes in kinesin-1, with full rotational freedom, and in response to changes in ionic strength, mutations, and the presence of microtubules. This has allowed us to uncover the inherent flexibility in kinesin-1 which gives insights into autoinhibition and the regulation of intracellular transport.

## INTRODUCTION

Kinesin-1, a ubiquitous molecular motor, plays a critical role in intracellular transport by moving many different kinds of cargo long distances along microtubule filaments. This diversity of cargos includes not only many different membrane bound organelles ^1–6^, but also RNA ^7–9^, cytosolic protein complexes ^10,11^ and microtubules themselves ^12,13^. While significant progress has been made towards understanding the fundamental mechanisms of how kinesin-1 turns ATP hydrolysis into motility ^14–16^, the regulation of its activity and ability to interact with such a diverse range of cargo in many different local cellular environments is not fully understood. Recent studies have highlighted the importance of conformational changes in modulating kinesin-1 function ^17–19^. However, the complex interplay between conformational changes and microtubule interactions requires further investigation.

The core unit of kinesin-1 is a homodimer of two heavy chains, KIF5A, B or C. These consist of the well conserved N-terminal motor domains, a long elongated coiled-coil stalk and an intrinsically disordered C-terminal tail region ^14,20^. Kinesin-1 homodimers can be supplemented by a pair of kinesin light chains (KLCs 1-4), but light chain independent functions of kinesin-1 are known ^2,8,12^ and several studies indicate that kinesin light chains are not required for motility *in vitro* ^21–23^. Indeed, both light chain bound and unbound populations can be purified from the brain ^24^ and protein copy number estimates indicate a consistent molar excess of heavy chains compared to light chains ^25^. Consequently, the kinesin-1 heavy chains alone are sufficient to recapitulate both autoinhibition and motility.

Basic ‘open’ and ‘closed’ conformational states of the kinesin-1 have been defined almost since its discovery ^26,27^. Kinesin-1 is a microtubule-stimulated ATPase ^28^, where autoinhibition prevents futile ATP consumption when kinesin is not bound to cargo. The tail of kinesin contains a conserved IAK motif that is important for autoinhibition via a “head to tail” interaction where the IAK motif docks between the motor domains ^29–32^; thus the ‘closed’ conformation of kinesin has became synonymous with an autoinhibited molecule. However, recent evidence suggests the IAK motif is not the sole component of autoinhibition, and other interactions may also be in play ^18,33^. By definition, autoinhibited conformations must be highly unfavourable when kinesin-1 is actively walking along microtubules, and so walking kinesin is synonymous with an ‘open’ conformation. Beyond this, the conformational landscape of kinesin-1 is poorly defined, including the conformational changes required to allow walking along microtubules and whether the ‘open’ conformation is a single conformer or represents a broader range of related structures. A binary open/closed model implies a rapid interconversion between these two states, although it is currently unknown if this is the case. Part of the reason that the conformational landscape of kinesin-1 remains poorly defined is likely the paucity of high resolution structural data for kinesin-1 beyond the well characterised motor domains. This in turn is compounded by commonly used methods in the field, such as singlemolecule walking assays, giving information only on the end product of these complex events as kinesin walks along microtubules. Consequently, the conformational changes within the stalk and tail of kinesin are often assumed rather than measured directly.

In order to better understand the ability of kinesin-1 to carry out such a diverse range of transport functions, we sought to establish an experimental pipeline that could capture the kinesin-1 conformational landscape. We wanted to be able to study freely diffusing molecules, as immobilisation of kinesin can cause aberrant activation through nonspecific interactions with the kinesin tail (a property leveraged for some motility studies of full length kinesin ^33,34^). We also wanted the ability to study kinesin in response to changes in both external and internal conditions; external conditions include factors such as ionic strength and pH, but also the presence and absence of microtubules and cargos, whilst the internal state of the protein includes mutations or posttranslational modifications. To this end we applied both single molecule Fluorescence Resonance Energy Transfer (smFRET) and Flow Induced Dispersion Analysis (FIDA) to the study of kinesin-1. smFRET in particular is emerging as a powerful tool for monitoring conformational changes, as the single molecule resolution provides information on multiple conformers, rather than averaged ensemble measurements of populations ^35,36^.

By utilizing self-labeling enzymes and cell-permeable fluorophores, we were able to probe the conformational landscape of kinesin-1 and its response to different external (ionic strength, microtubules) and internal (mutants) conditions. Given that dimers of kinesin-1 heavy chains recapitulate the two key features of kinesin, motility and autoinhibition, we reasoned that key conformational changes could be captured with heavy chains alone. Our findings reveal the dynamic conformational landscape of kinesin-1 dimers, with ionic strength and microtubule binding playing crucial roles in modulating the equilibrium between open and closed states. These insights provide a deeper understanding of the molecular mechanisms underlying kinesin-1 function and have implications for understanding the regulation of intracellular transport.

## RESULTS

### Self-labelling enzymes and cell permeable fluorophores are suitable for smFRET

To measure conformational changes in kinesin-1, we chose to use the self labelling enzymes CLIP and SNAP to position fluorophores at the N- and C-termini of kinesin-1. Although dependent on the fluorophores in use, the typical dynamic range for FRET occurs when fluorophores are 3-10 nm apart. The extended conformation of kinesin is thought to be between 60 and 80 nm ^26,37,38^, greatly exceeding this limit, whilst the autoinhibited conformation should bring the N-& C-termini into close proximity ^29,30^ creating a signal change upon closing of kinesin ^39^. As of yet there are no experimentally determined high resolution structures of the kinesin-1 stalk region that would allow the considered positioning of fluorophores within the coiled coils by direct labelling approaches. Consequently, self labelling enzymes allow the conjugation of bright fluorophores to proteins, without the additional challenges imposed by the maturation of fluorescent proteins and their concomitant spectra ^40^. We chose to pair SNAP and CLIP, as they are matched in size (19.4 kDa) and are smaller than other self labelling enzymes such as the HaloTag (33 kDa); as FRET is highly distance dependent, this is a key advantage.

To test whether the CLIP tag and SNAP tag had the capacity to be used for smFRET experiments with readily available fluorescent ligands, both were fused with a short flexible linker to provide a high FRET positive control (CLIP-SNAP, Figure 1A). HEK 293 cells were transfected with CLIP-SNAP and labelled with the cell permeable fluorophores CLIP-Cell TMR-Star and SNAP-Cell 647-SiR (Figure 1B), the cell lysate was collected and used directly for measurements. High quality confocal smFRET measurements rely on the diffusion of a single molecule through the confocal volume at any one time (Figure 1C). In practice, although events are short (a few milliseconds), an observation rate of about one molecule per second avoids coincidence events of two or more molecules in the volume at a time producing erroneous FRET measurements (Figure 1D). Any unbound ligand or other background contaminants contribute to this rate of observation, increasing acquisition time to many hours and making experiments impractical. Our preliminary results were promising and FRET bursts were observed, but the background of unbound ligands made acquisition very slow.

**Figure 1:**
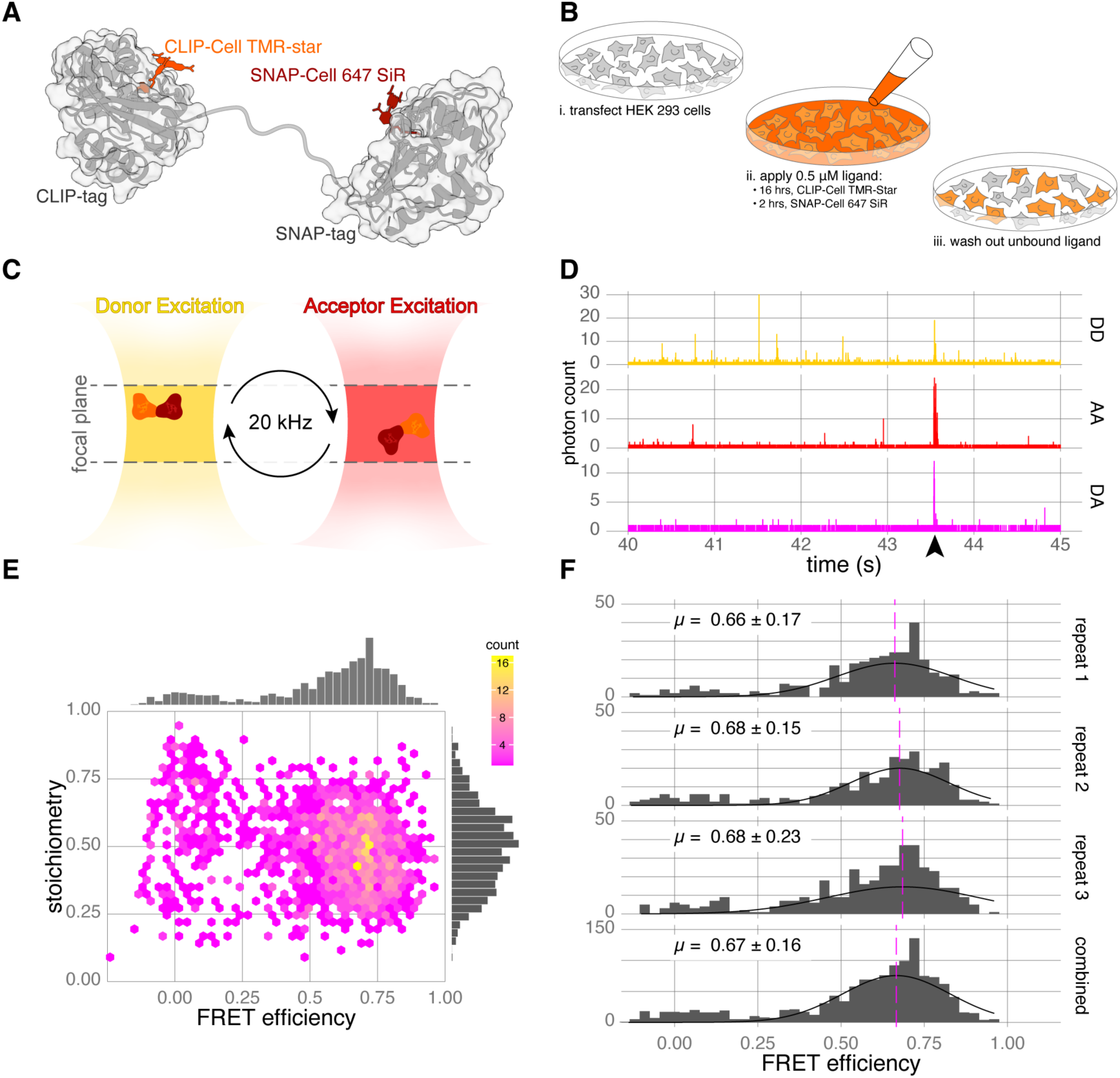
A CLIP-SNAP fusion protein is suitable for smFRET measurements using live-cell compatible fluorescent ligands. (A) Schematic of the labelled CLIP-SNAP fusion protein with the flexible linker (based on PDB: 6Y8P). (B) Steps for sample preparation requiring optimisation to minimise background contribution from unbound ligand. (C) smFRET in the confocal volume using alternating laser excitation (ALEX); only one molecule is present in the volume at any one time. Single molecules diffuse through the confocal volume (∼1 m^3^) whilst being probed by ALEX cycles of donor and acceptor excitation at 20 kHz ^50^. (D) Example time traces of TMR-Star and 647-SiR labelled CLIP-SNAP from the donor:donor (DD), acceptor:acceptor (AA) and donor:acceptor (DA) excitation:emission channels. FRET is characterised by a photon burst in all three channels (marked by arrowhead). (E) Hexbin plot of FRET efficiency versus stoichiometry for the CLIP-SNAP fusion protein expressed in HEK 293 cells. Data from five biological replicates, n = 1389 bursts. (F) Histograms of FRET efficiency from three biological repeats demonstrating reproducibility of live cell labelling for smFRET. Stoichiometry is gated to 0.25 and 0.75.

To eliminate background contributions from unbound fluorescent ligands, we undertook a systematic optimization of labelling for CLIP and SNAP tagged protein expressed in HEK 293 cells, and the cell permeable fluorophores CLIP-Cell TMR-Star and SNAP-Cell 647-SiR. This included using flow cytometry to calculate the optimal washout time for unbound ligand using untransfected cells (Figures S1A to S1B), and in-gel fluorescence of transfected cell lysate to optimise both ligand incubation time (Figures S1C to S1D) and concentration (Figures S1E to S1F). As previously shown ^41^, the labelling kinetics for CLIP tag are slower than SNAP tag, and our final labelling parameters were 0.5 *µ*M of CLIP-Cell TMR-Star and SNAP-Cell 647 SiR ligands on cells for 16 hours and 2 hours respectively, followed by 2 hours in ligand-free media to wash out unbound fluorophores. We observed minimal cross reactivity of labelling between CLIP and SNAP tag with these conditions (Figure S1G). Expressing and labelling protein in this way circumvented the need for protein purification, taking advantage of the cell membranes to act as dialysis membranes and remove unbound ligands (Figure 1B). Using in-gel fluorescence, we carried out similar experiments with the HaloTag and the TMR ligand, which showed some non-specific binding on the same time scales (Figure S1H).

Having removed contamination from the sample, the assay buffer became our biggest contributor to background - specifically BSA, regardless of source (Figure S1I). BSA is frequently added to *in vitro* assays of kinesin function as a stabiliser. We devised an in-house method to pre-bleach the BSA (see Materials and Methods, Figure S1J), which eliminated this final source of contamination. However, it is still the case that any use of coloured plastics in protocols (tips, tubes etc) or other significant coloured items (e.g. hair dye) is sufficient to contaminate reagents and render data acquisition impossible. Following optimization, background photon counts of diluted untransfected cell lysates derived from cells exposed to the ligand labelling procedure were indistinguishable from the assay buffer alone (Figures S1K to S1L).

With our optimised labelling conditions we observed robust smFRET signals of the CLIP-SNAP fusion protein after expression and dilution of lysate harvested from transfected HEK 293 cells. Each labelled molecule diffusing through the confocal volume results in a burst of photons (Figure 1D). Photons can be separated into three channels based on their excitation and emission. These are: donor-only (DD), acceptor-only (AA) and donor-acceptor (DA), also known as the FRET channel (Figure 1D). The number of photons in each of these channels is used to calculate the FRET efficiency (a measure of energy transfer from donor to acceptor fluorophore, and is inversely proportional to the distance between them) and stoichiometry for that molecule (a ratio of donor to acceptor labelling). Plotting FRET efficiency vs stoichiometry as a frequency for all observed molecules of the CLIP-SNAP fusion protein (Figure 1E), shows a peak at 0.68 and 0.49 for FRET efficiency and stoichiometry respectively. Further, estimates of mean FRET efficiency (*±* standard deviation) for three independent biological replicates were 0.66 *±* 0.17, 0.68 *±* 0.15 and 0.68 *±* 0.23 (Figure 1F), demonstrating a high degree of reproducibility. Having established that CLIP tag and SNAP tag together with cell permeable fluorophores are suitable for smFRET, we applied our labelling method to kinesin-1.

### HEK 293 cells are a suitable expression system for full length motor protein studies

We chose to focus the majority of our analysis on KIF5A (figures 2A to 2B). The core of the mammalian kinesin-1 motor is a homodimer of two heavy chains, either KIF5A, KIF5B or KIF5C. As with many protein complexes, AlphaFold has been able to provide the first structural predictions of the full kinesin-1 homodimer (Figure 2B and Figure S2A). These largely match previous predictions for the motor domain, coiled-coil stalk and intrinsically disordered regions ^42,43^. However, there is one notable difference: previous models suggested the major hinge that brought the head and tail of kinesin together was between CC1 and CC2^34,44^. This unstructured region is now predicted to form a retrograde loop (residues V565 to A579) that facilitates the interlocking of these coiled coils together. It is also the case that structural predictions of the kinesin-1 stalk have changed with different versions of AlphaFold. The extended conformation of the tail that was reliably produced by earlier versions of AlphaFold (Figure S2B as example), has now become more compact, with coiled-coils stacked against each other to create a concertina structure (Figure S2C). The latest version, AlphaFold3, still produces a concertina structure when the tail is modelled, although we noticed that hydrogen bonding compacting the coiled coils is even more extensive. Within the concertina, flexible regions between CC2/3 and CC3/4 define two new hinge regions, hinge 1 & 2 (residues K685-T689 and M815-G821 respectively, Figure 2B), providing potential for a large range of movement in the kinesin stalk and tail.

**Figure 2:**
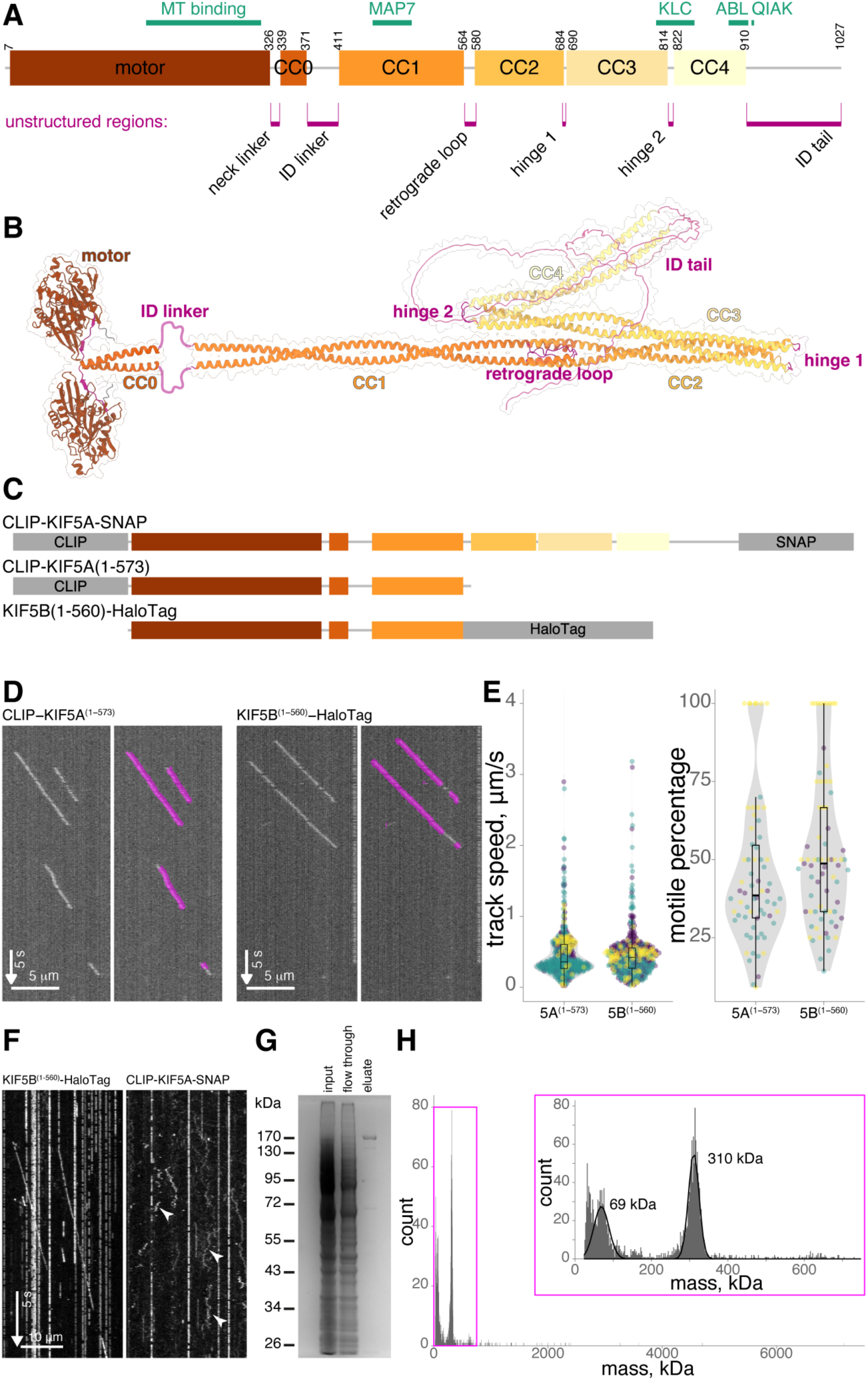
Tagged KIF5A is functional and suitable for smFRET assessment. (A) Schematic representation of the domain structure of mouse KIF5A, including motor domain and coiled-coil (CC) regions 0-4. Key binding motifs are illustrated in green and unstructured regions in pink. (B) AlphaFold prediction of dimeric KIF5A, coloured as domain structure above. Regions 1-371 and 411-1027 were modelled separately to show the elongated form. (C) Domain schematics illustrating CLIP, SNAP and HaloTag positions in respective constructs. (D) Example kymographs of KIF5B(1-560)-HaloTag and CLIP-KIF5A(1-573) from single molecule walking experiments (left) and overlaid with tracking data in magenta (right). (E) Analysed particles from experiments in (C) showed no significant difference in track speed (n = 401 and 414 particles for CLIP-KIF5A(1-573) and KIF5B(1-560)-HaloTag respectively, p = 0.7 by Mann-Whitney U) or motile percentage (n = 71 and n = 66 microtubules for KIF5B(1-560)-HaloTag and CLIP-KIF5A(1-573) respectively). Data from three biological replicates. (F) Single molecule walking experiments of KIF5B(1-560)-HaloTag compared to CLIP-KIF5A-SNAP show intact autoinhibition of the latter. Arrowheads highlight non processive binding events. (G) Coomassie stained SDS-PAGE gel showing pull down and elution of CLIP-KIF5A-SNAP using the incorporated FLAG tag. (H) Mass photometry of eluate in G with a 310 kDa peak corresponding to the tagged KIF5A dimer.

Having established CLIP and SNAP tags as suitable for smFRET, we moved to applying this labelling strategy to kinesin-1 by positioning CLIP and SNAP at the N- and C-termini of KIF5A (CLIP-KIF5A-SNAP, Figure 2C). However, before proceeding to smFRET analysis, we wanted to ensure that tagging kinesin at both the N and C termini simultaneously does not interfere with KIF5A motility or autoinhibition. *In vitro* reconstitution using cell lysates as a source of kinesin protein, is a well established technique for assessing kinesin motility on isolated microtubules ^18,21,45^. Using single molecule total internal reflection fluorescence microscopy (TIRFM) assays (Figure 2D), comparing the motility of the constitutively active truncation CLIP-KIF5A(1-573) to well studied C-terminally tagged KIF5B(1-560)-HaloTag, showed no difference in track speed or motile percentage (Figure 2E). Autoinhibition of full length kinesin is also conserved in this assay system, with many fewer events typically observed compared to constitutively active mutants ^18^. Similarly, in comparison to KIF5B(1-560)-HaloTag, we observed robust autoinhibition of CLIP-KIF5A-SNAP in TIRFM assays (Figure 2F). Although labelled CLIP-KIF5A-SNAP can be observed interacting with the microtubules (Figure 2F, arrowheads), the particle movement is diffusive across the lattice rather than the unidirectional processive motility observed for active motors. We conclude that motility and autoinhibition are conserved in CLIP-KIF5A-SNAP.

As a neuron specific isoform of kinesin-1, KIF5A is not expressed endogenously in our HEK 293 expression system, so co-purification with untagged endogenous kinesin-1 subunits is unlikely. To confirm that no associations of this type were interfering with our analysis, we performed both co-immunoprecipitation (Figure S2D) and mass photometry (figures 2G to 2H). When HEK 293 cells were transfected with CLIP-KIF5A-SNAP (or CLIP-KIF5B-SNAP), we were never able to observe a co-immunoprecipitation with endogenous KIF5B (Figure S2D). To carry out a mass photometry analysis of complexes, we took advantage of the C-terminal FLAG included in our CLIP-SNAP expression vector (pCLAP). Complexes were enriched on FLAG-trap beads and eluted with the FLAG peptide prior to analysis (Figure 2G). The largest molecular weight complex observed with this method had a molecular weight of ∼310 kDa, corresponding well to the predicted size of a homodimer of CLIP-KIF5A-SNAP (Figure 2H). A consistent association with kinesin light chains would be expected to produce a peak at ∼450 kDa, but no peak is evident in this region. Likewise, no monomers of KIF5A were observed, indicating the homogeneity of the preparation. Having now established our expression system, labelling and assay procedures, we undertook an smFRET analysis of kinesin-1.

### Relative labelling efficiencies of SNAP and CLIP give dimeric molecules a unique stoichiometry signature in smFRET

Based on current models of kinesin-1 regulation, we anticipated that the majority of CLIP-KIF5A-SNAP should be in a closed conformation. It was also possible that we would observe both the open (low FRET) and closed (high FRET) conformations, as in recent negative stain electron microscopy (EM) studies of purified kinesin ^17,18^. Plotting FRET efficiency versus stoichiometry for CLIP-KIF5A-SNAP FRET bursts shows a single continuous population with a long tail into the higher FRET regions of the axis (Figure 3A). This does not support a model of rapid transition between two distinct and stable conformational states (median FRET efficiency is 0.136). However, compared to the CLIP-SNAP control protein, we also noticed a drop in stoichiometry; CLIP-SNAP stoichiometry is 0.49 *±* 0.12, whilst CLIP-KIF5A-SNAP is 0.39 *±* 0.17 (mean *±* sd). Given that labelling kinetics are slower for CLIP tag compared to SNAP tag ^41^, this shift is most likely to have arisen from the difference in labelling efficiency between SNAP and CLIP for dimeric kinesin-1 (Figure 3B). This would result in a subpopulation of CLIP-KIF5A-SNAP dimers that carry two SNAP ligands for every CLIP ligand.

**Figure 3:**
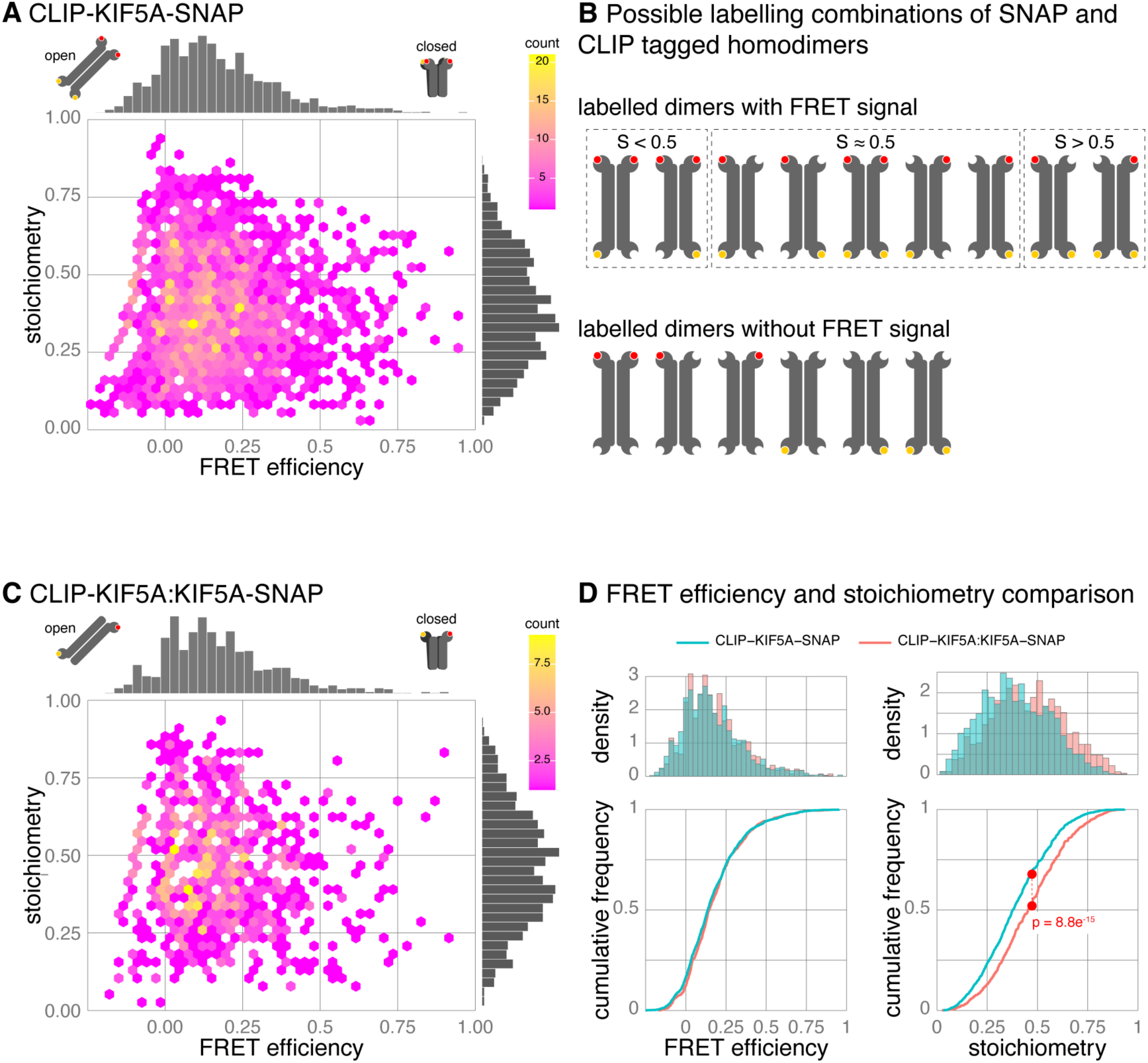
Tagging strategy affects stoichiometry but not FRET efficiency. (A) Hexbin plot of FRET efficiency versus stoichiometry for CLIP-KIF5A-SNAP. Data combined from five biological replicates, n = 2436 bursts. (B) Schematic of possible labelling stoichiometries of CLIP-SNAP tagged homodimers and the corresponding stoichiometry value. Labelled CLIP and SNAP are yellow and red circles respectively. (C) Hexbin plot of FRET efficiency versus stoichiometry for heterodimers formed by CLIP-KIF5A:KIF5A-SNAP. Data from three biological replicates, n = 901 bursts. (D) Histograms and cumulative frequency plots of data from (A) and (C) comparing FRET efficiency & stoichiometry of KIF5A dimers formed of CLIP-KIF5A-SNAP (blue) or CLIP-KIF5A:KIF5A-SNAP (red).

To test if labelling efficiency impacts stoichiometry and ensure this in turn does not affect our interpretation of the FRET efficiency distribution, we created KIF5A dimers, by co-transfection of KIF5A tagged with a single CLIP tag or SNAP tag (CLIP-KIF5A:KIF5A-SNAP). This strategy ensures any dual labelled molecules can only have one CLIP and one SNAP tag, ensuring equivalent labelling and a stoichiometry of 0.5. CLIP-KIF5A:KIF5A-SNAP dimers were labelled and analysed with our optimised protocol. Plotting FRET efficiency versus stoichiometry of CLIP-KIF5A:KIF5A-SNAP shows the predicted increase in stoichiometry with mean *±* sd of 0.46 *±* 0.18 (figures 3C to 3D). Further, comparing the stoichiometry of biological replicates for the two labelling regimes highlights the consistency in stoichiometry for CLIP-KIF5A:KIF5A-SNAP compared to a more variable response for CLIP-KIF5A-SNAP dimers (Figure S3A). However, despite this impact on stoichiometry, the FRET efficiency distribution between the two labelling regimes was almost identical (Figure 3D), maintaining the skewed normal distribution with a long tail into the higher FRET regions of the axis. One important side effect of the single labelling approach is having to rely on the dimerization of CLIP-KIF5A:KIF5A-SNAP, accounting for (at most) 25% of the KIF5A dimers assembled. In practice the percentage of dual labelled kinesins is likely much lower for two reasons: first, not all cells are co-transfected, and; second, data processing of smFRET photon bursts selects for molecules with both donor and acceptor fluorophore emission, losing molecules via inefficient CLIP labelling. Consequently, our ability to accumulate dual labelled bursts from the single tagging approach was severely impacted, greatly increasing the time for acquisition. A typical acquisition per biological replicate is three 15 minute acquisition periods of the same sample, per construct (approximately an hour of instrument time); a FRET burst frequency at a quarter of the rate requires at least four hours of acquisition. These long time periods at room temperature can potentially lead to sample integrity issues, particularly when multiple constructs are being compared within an experiment. Given this, combined with minimal impact on FRET efficiency distribution, we chose to use double tagged kinesin constructs for our studies going forward.

### Changing the ionic environment alters the KIF5A conformational landscape

Given the long continuous FRET efficiency distribution we observed in our first experiments with tagged KIF5A (Figure 3), indicative of continuous sampling of many conformational states, we next sought to capture isolated populations of open or closed conformations.

Altering the ionic strength of the assay buffer has been used for many years as a strategy for altering the conformation of kinesin-1. Original studies were able to indicate a transition from a compact state of kinesin to an elongated state by both rotary shadowed EM ^26^, and changing sedimentation coefficients (a proxy for hydrodynamic radius) ^27,46^, in response to increasing ionic strength up to an additional 1 M. Consequently, we predicted that the addition of NaCl to our assay buffer would alter the FRET efficiency distribution.

To study the impact of increasing ionic strength on kinesin conformation, HEK 293 cells were co-transfected with CLIP-KIF5A-SNAP, and labelled with our optimised protocol. The same kinesin sample was then diluted either into smFRET assay buffer, which has low ionic strength, or assay buffer supplemented with an additional 150 mM or 800 mM NaCl, immediately prior to data collection. Accordingly, stoichiometry was consistent across samples (Figures S4A to S4C). However, changes in FRET efficiency were apparent. Compared to the no additional NaCl condition, adding 150 mM NaCl caused a significant increase in the FRET efficiency distribution, clearly visible in a shift to the right in the cumulative frequency distribution (figures 4A to 4B). Adding 800 mM NaCl also caused an increase, but the shift was less pronounced. In fact, we found the response to 800 mM NaCl highly variable, often losing measurable kinesin molecules from the sample due to salting-out and precipitation, most obvious when comparing FRET efficiency across biological replicates (Figure 4C).

**Figure 4:**
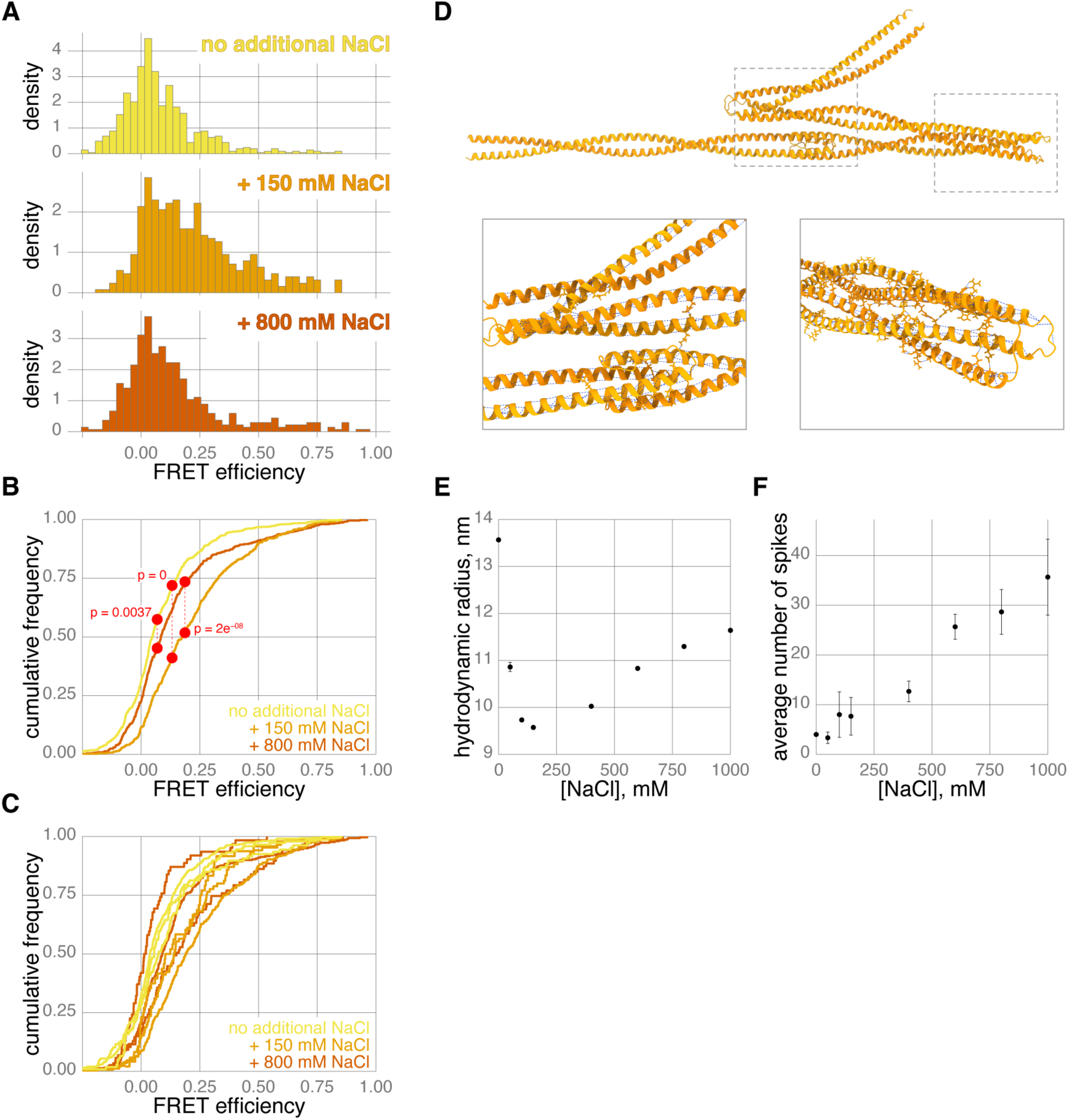
Changing salt concentration changes KIF5A behaviour. (A) Histograms of FRET efficiency for CLIP-KIF5A-SNAP in changing NaCl concentrations. Data combined from three biological replicates, n = 677, 421 and 449 bursts for 0 (yellow), 150 (orange) and 800 mM (brown) additional NaCl respectively. (B) Cumulative frequency plots of data in (A) comparing FRET efficiency for CLIP-KIF5A-SNAP in changing NaCl concentrations. (C) Cumulative frequency plots comparing FRET efficiency for individual biological replicates of CLIP-KIF5A-SNAP in changing NaCl concentrations. (D) AlphaFold prediction of KIF5A(411-911) with expanded regions highlighting alpha-helix packing and hydrogen bonding both where CC domains 1-4 stack and the hairpin formed between CC2 and CC3. (E) Apparent hydrodynamic radius of CLIP-KIF5A-SNAP for increasing additional NaCl concentrations determined by FIDA. Data points are mean *±* standard deviation of three replicates. (F) Spikes, corresponding to aggregated protein, observed in analysed Taylorgrams for increasing additional NaCl concentrations. Data points are mean *±* standard deviation of three replicates.

If the increase in FRET efficiency at 150 mM NaCl represents an increase in compact folded kinesin motors, this is a highly unexpected result, as raising the ionic concentration is more typically linked to the disruption of protein-protein interactions. However, this observation could be explained by disruption of the hydrogen bonding and electrostatic interactions holding the concertina structure of the kinesin-1 stalk (Figure 2B) in place. Hydrogen bonding across CC1-4 is a consistent factor positioning stacking of coiled-coils in AlphaFold predictions of the kinesin stalk. Although the exact number and positions of interactions across the stacked coiled-coils varied, their presence was consistent across AlphaFold predictions of the kinesin-1 stalks of mammalian isoforms (Figure 4D & Figures S4D to S4E) using AlphaFold 2.3 onwards. Disruption of the hydrogen bonding and electrostatic interactions holding the concertina in place could directly facilitate a closer association of the kinesin head and tail, resulting in a more compact conformation observed as an increase in the FRET efficiency.

A large change in conformation from an elongated to a compact configuration should also be observable as a change in the hydrodynamic radius (Rh). To test this hypothesis, we performed an independent assay to analyse the hydrodynamic radius of kinesin prepared by binding and elution from FLAG beads followed by Flow Induced Dispersion Analysis (FIDA, Figure 4E, Figure 4F & Figure S4F). In contrast to confocal smFRET measurements which are in the femto-molar range with single molecules observed for ∼2 ms, FIDA measurements are carried out in the nano-molar range, giving an ensemble measurement over a ∼2 minute period. In strong agreement to the smFRET measurements of kinesin in diluted cell lysate, FLAG purified CLIP-KIF5A-SNAP was also in the most extended conformation in the absence of additional NaCl (mean Rh is 13.6 *±* 0.03), reaching its most compact state with 150 mM NaCl (mean Rh is 9.57 *±* 0.02). Similar to smFRET results, an extension at higher NaCl concentrations can also be observed (mean Rh at 800 mM is 11.30 *±* 0.04), though this is still more compact than the conformation with no additional NaCl. Globular proteins demonstrate a proportional relationship between their molecular weight and their hydrodynamic radii, which becomes distorted by extended conformations. The molecular weight of a CLIP-KIF5A-SNAP dimer is ∼318 kDa, which if purely globular would give a hydrodynamic radius of 6.59 nm. However, the experimentally derived Rh of 13.6 nm, corresponds to a molecular weight of 2.2 MDa for a globular protein, almost seven times the mass of a CLIP-KIF5A-SNAP dimer. Even at 150 mM NaCl kinesin-1 is still relatively elongated, as Rh of 9.57 nm corresponds to a molecular weight of ∼850 kDa for a globular protein. Another feature we noticed by FIDA was the increasing number of spikes in the signal with increasing NaCl concentration (Figure 4F and Figure S4F), corresponding to protein aggregates passing through the capillary ^47^. This confirmed our observations from smFRET, that kinesin-1 was likely salting out at high NaCl concentrations (Figure 4C). Taken together, our results from FIDA and smFRET show excellent agreement overall, with the lowest FRET occurring in conditions with the highest hydrodynamic radius and vice versa.

### An activating mutation of KIF5A does not increase the frequency of the open conformation

KIF5A has an extended disordered tail (Figure 2B) in comparison to KIF5B and KIF5C (106 compared to 40 and 32 residues respectively), which could easily reduce any FRET efficiency measurements irrespective of the conformation of the coiled-coil stalk. To ensure this was not the case and that results are consistent across KIF5 isoforms, we analysed a mutant KIF5A with the disordered region truncated to the same length as KIF5B. When analysed by smFRET, CLIP-KIF5A(1-953)-SNAP shared the same key characteristics of the full length KIF5A (Figure 5A, Figure 5B & Figure 5E), namely one continuous skewed distribution which includes high FRET events, but a median FRET efficiency of 0.116. As a direct comparison we also performed smFRET analysis on CLIP-KIF5B-SNAP with similar results (Figure 5C, median FRET efficiency of 0.143). However, it was noticeable that KIF5B constructs were consistently more difficult to overexpress in HEK 293 cells (Figure S5A), potentially due to intact degradation pathways and regulation of expression levels of an endogenous subunit, resulting in difficulty accumulating bursts. When comparing the cumulative frequency plots of KIF5B to either full length KIF5A or KIF5A(1-953), there was a subtle increase in FRET efficiency for KIF5B (Figure S5B and Figure S5C respectively), however the overall distribution was broadly similar.

**Figure 5:**
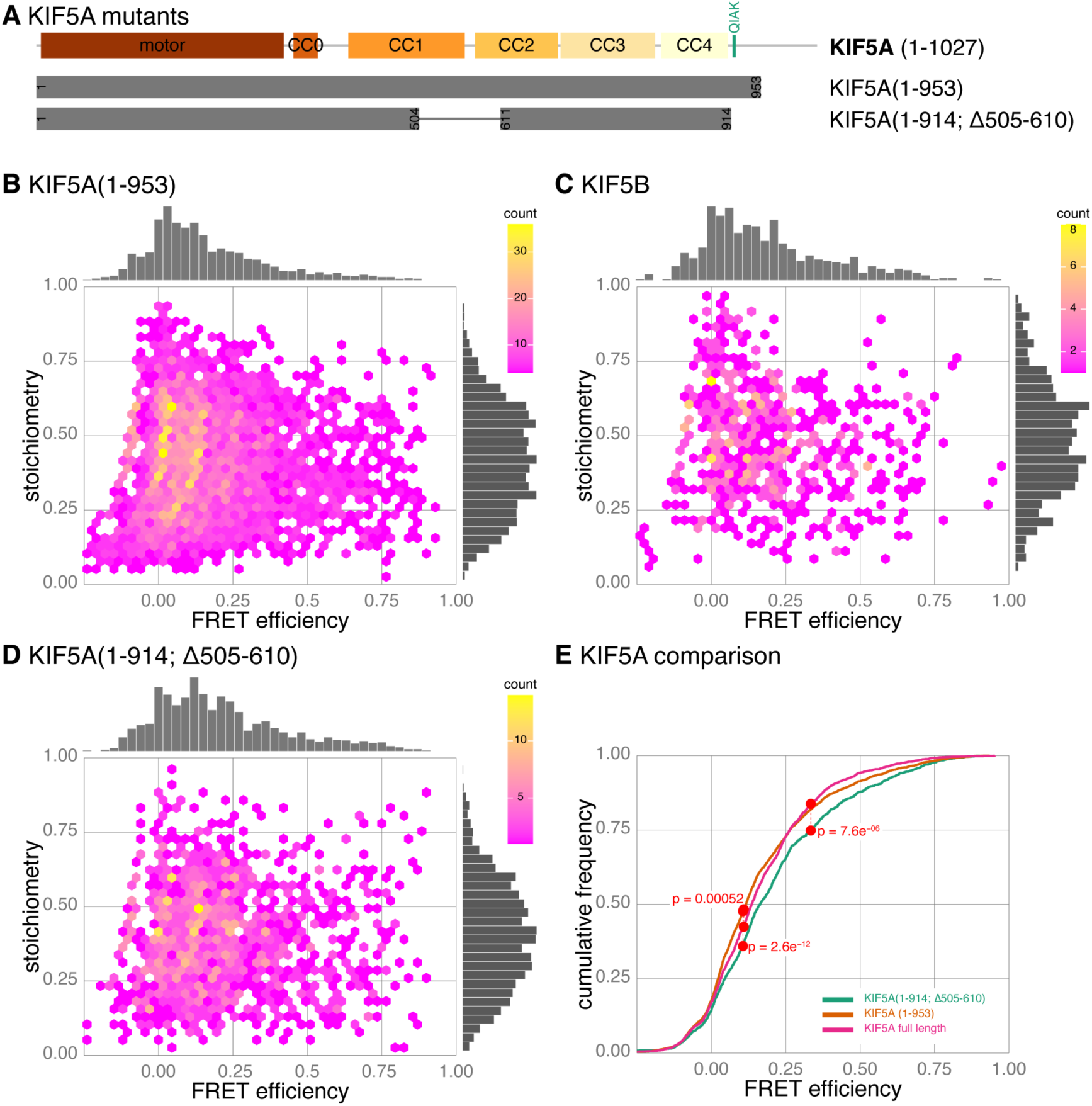
Activating mutations of KIF5A alter FRET efficiency. (A) Schematic of KIF5A mutants used in relation to the domain structure of KIF5A. (B - D) Hexbin plots of FRET efficiency versus stoichiometry for CLIP-KIF5A(1-953)-SNAP (B), CLIP-KIF5B-SNAP (C) and CLIP-KIF5A(1-914; Δ505-610)-SNAP (D). Data combined from four to six biological replicates, n = 3800, 666 and 1475 bursts respectively. (E) Cumulative frequency plots of data in (B & D) comparing FRET efficiencies for KIF5A(1-953) and KIF5A(1-914; Δ505-610) mutants to full length KIF5A.

To further investigate the conformational landscape of kinesin-1, we created a double mutant that should retain much of the length of the motor whilst reducing autoinhibition (Figure 5A). In one of the key studies that revealed autoinhibition of kinesin-1, Friedman and Vale pinpointed residues 505-610 of human KIF5B as a potential hinge region facilitating the head-tail interaction of kinesin for autoinhibition; by deleting this region both the velocity and event frequency of *in vitro* walking events was increased, ^44^, an observation that was recently repeated ^18^. The structure of this region, which includes the retrograde loop, (highlighted in Figure S5D) is currently not predicted to be flexible by AlphaFold, but high resolution structural data to confirm this is still lacking. In order to further open up the motor, we also deleted residues 915-1027 of the tail which includes the IAK motif, removing the head-tail interaction and creating a constitutively active motor ^18^. Consequently, we predicted that this mutant would show a decrease in FRET efficiency as it was far more likely to be in the open conformation. In fact, CLIP-KIF5A(1-914; Δ505-610)-SNAP showed an increase in FRET efficiency (figures 5D to 5E; median FRET efficiency of 0.164). One possible explanation is that removing residues 505-610 (whilst retaining the coiled-coil structure through the heptad repeat, Figure S5E), disrupts the predicted concertina structure of the stalk increasing the likelihood of the head and tail coming into proximity despite a lack of interaction between motor domains and IAK motif.

### Kinesin-1 is highly flexible, but crosslinking differentiates between activating kinesin mutants and intact autoinhibition

The long continuous distributions we observe by smFRET are suggestive of a broad conformational landscape and extensive flexibility in kinesin-1. To investigate the basis of this flexibility further, we calculated AlphaFold predictions of individual stalk domains (CC1-2, CC3 and CC4) in isolation across mammalian isoforms (Figure S6A). The predicted stalk-domain structures have high confidence pLDDT scores for the majority of their length. However it was noticeable that the phase of the predicted coiled-coil structures breaks down at the interface between these domains, accompanied by a decrease in pLDDT score (Figure S6A, grey boxes). This means that the phase of each individual coiled-coil domain is often out of sync from one coiled coil to the next, likely providing structural tension at the flexible regions of hinge 1 and hinge 2 (Figure 2B). Consequently, when CC1-4 is modelled as one complete stalk domain the overall pLDDT confidence score drops, particularly around the hinges, a strong indicator of conformational flexibility in these regions (Figure S6B).

Based on the observed skewed, continuous, FRET efficiency distributions of kinesin-1, we hypothesise that kinesin-1 is sampling a large conformational space in solution, transiently bringing head and tail into closer proximity only some of the time. Recent negative stain EM studies of kinesin-1 have made extensive use of crosslinking reagents to stabilise closed conformations after large scale protein purification ^17–19^. In our kinesin preparations, we have not enriched for a specific conformational state by size exclusion chromatography. However, we reasoned that crosslinking our full length kinesin-1 would still capture some of these transient closed conformations, stabilising them enough to increase FRET efficiency.

To carry out crosslinking, labelled CLIP-KIF5A-SNAP was enriched from cell lysates with FLAG-trap beads, eluted with FLAG peptide and then subjected to crosslinking with a 1:1 ratio by mass with the amine-to-amine crosslinker bis(sulfosuccinimidyl)suberate, BS3. The BS3 ratio was determined to be optimal for preserving dimers without the over crosslinking observed at higher ratios (Figures S6C to S6E). We did observe an impact of crosslinking on stoichiometry within these experiments (Figure S6F; mean stoichiometry of 0.46 and 0.51, for uncrosslinked and crosslinked samples, respectively) likely due to stochastic disruption of the fluorophores by BS3. However, based on previous observations (Figure 3D) we do not think this negatively impacts our FRET efficiency observations (see below).

Compared to uncrosslinked kinesin from the same preparation, we observed a small but consistent increase in FRET efficiency for CLIP-KIF5A-SNAP (Figure 6A, Figure 6E; median FRET efficiency for uncrosslinked and crosslinked is 0.082 and 0.119, respectively), consistent with trapping of the head-to-tail interaction. A similar change is also observed for the shortened KIF5A that nevertheless preserves autoinhibition, CLIP-KIF5A(1-953)-SNAP (Figure 6B, Figure 6F; median FRET efficiency for uncrosslinked and crosslinked is 0.113 and 0.147, respectively). In further support of this model, CLIP-KIF5A(1-914; Δ505-610), the KIF5A construct where autoinhibition is disrupted, shows the opposite effect in response to crosslinking with BS3, a decrease in FRET efficiency (Figure 6C, Figure 6G; median FRET efficiency for uncrosslinked and crosslinked is 0.132 and 0.085, respectively), reflecting an increased tendency to occupy an open conformation.

**Figure 6:**
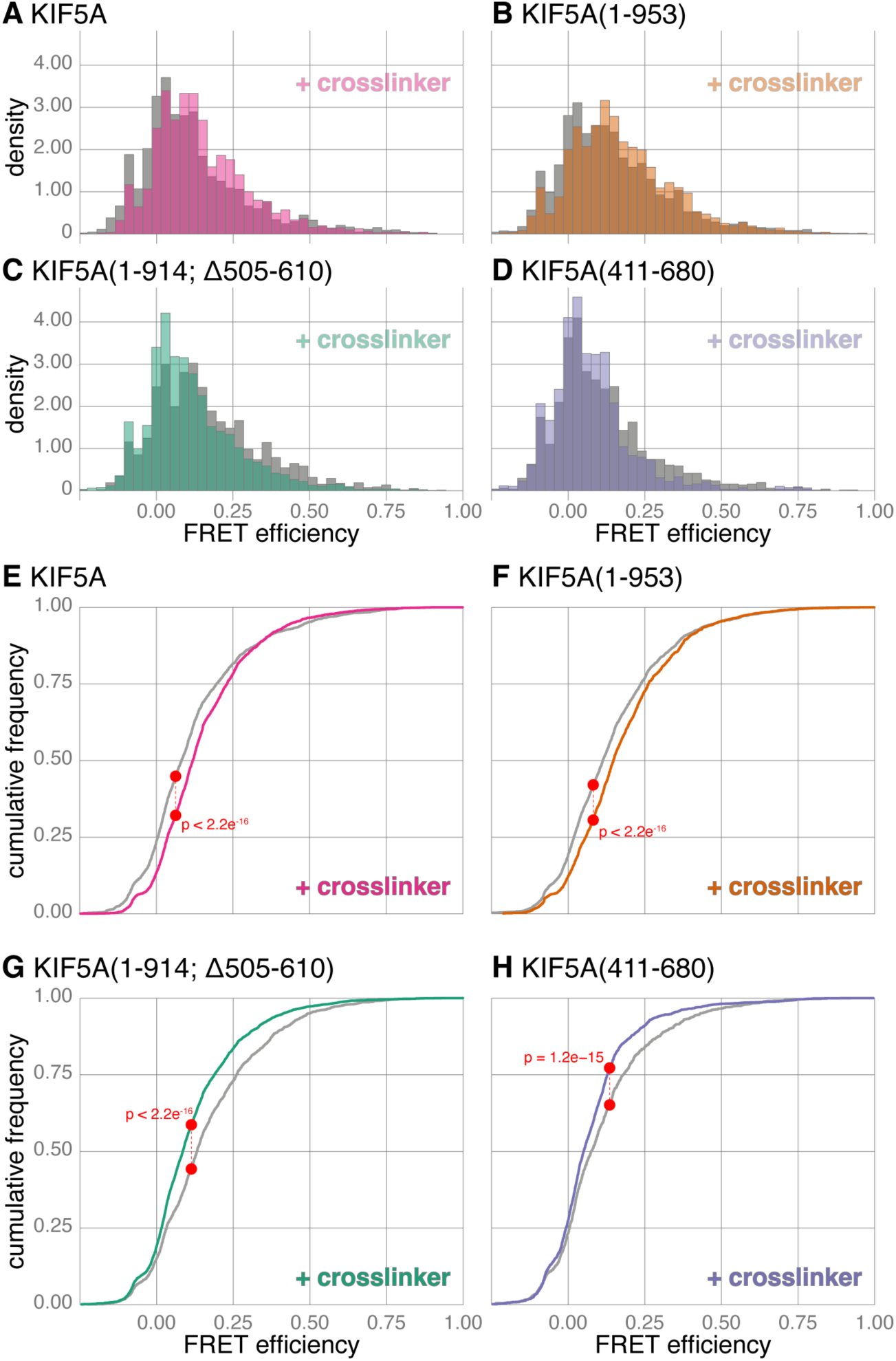
Crosslinking KIF5A constructs reveals a differential response to activating vs truncating mutations. (A - D) Histograms comparing the FRET efficiency of uncrosslinked (grey) and crosslinked (coloured overlay) protein samples of KIF5A (A), KIF5A(1-953) (B), KIF5A(1-914; Δ505-610) (C), and KIF5A(411-680) (D). Data combined from five biological replicates, and n bursts for crosslinked & uncrosslinked samples as follows: KIF5A, 4093 & 3066; KIF5A(1-953), 4031 & 2873; KIF5A(1-914; Δ505-610), 6002 & 2626; and KIF5A(411-680), 3925 & 1700. (E - H) Cumulative frequency plots of data in (A-D) comparing FRET efficiencies of uncrosslinked (grey) and crosslinked (coloured) protein samples of KIF5A (E), KIF5A(1-953) (F), KIF5A(1-914; Δ505-610) (G), and KIF5A(411-680) (H).

Alphafold predicted structures of the combined CC1-2 domain consistently have the highest pLDDT scores of any domain within the kinesin-1 stalk, whether modelled alone or in combination with other domains. To better understand whether this domain was contributing to the flexibility of the kinesin-1 motor, we carried out a smFRET analysis of CLIP-KIF5A(411-680)-SNAP with and without crosslinking. The CC1-2 domain is predicted by AlphaFold to be ∼35 nm long, and CLIP-KIF5A(411-680)-SNAP gave the lowest FRET efficiency of any construct we had observed to date (median FRET efficiency 0.075) consistent with the majority of fluorophores being outside the range amenable to a FRET interaction (10 nm). However, despite this low FRET efficiency, crosslinking CLIP-KIF5A(411-680)-SNAP was able to reduce the FRET efficiency even further (median FRET efficiency 0.051). This result indicates that cross-linking CLIP-KIF5A(411-680)-SNAP traps it in a more extended conformation, suggesting that even CC1-2 retains some flexibility along its length, and can therefore influence the conformational landscape of kinesin-1 (Figure 6D, Figure 6H).

### The presence of microtubules polarises the kinesin-1 conformational landscape observed by smFRET

All smFRET and FIDA observations of kinesin-1 thus far in this study have been carried out in the absence of microtubules. However, first and foremost kinesin-1 is a microtubule-stimulated ATPase and so its response to microtubules is an essential component for understanding how kinesin functions in an integrated system within cells.

A key benefit of using confocal smFRET to study intramolecular kinesin dynamics is that kinesin conformation can be observed with and without microtubules or cargo. We therefore used smFRET to ask how the dynamic conformational landscape of kinesin motors is affected by microtubules. We added unlabelled, taxol stabilised microtubules to our smFRET assay so that within the assay volume, kinesin-1 molecules were free to interact stochastically with the microtubules. However, unlike TIRF measurements, where kinesin observations are limited explicitly to those only interacting with the surface of the microtubule, confocal smFRET will always sample a mixed population of kinesins that may or may not be interacting with microtubules at the moment of observation. We reasoned that due to their larger size, kinesins bound to microtubules would diffuse more slowly through the confocal volume. We therefore compared photon bursts that were longer than 0.75 ms in duration, to improve our chances of capturing microtubule interacting kinesins. Previous cryo-EM studies have been successful in trapping autoinhibited kinesin-1 motor domains bound to microtubules ^29,32^. Intriguingly, when we used smFRET to look at the distribution of kinesin molecules within our slower diffusing kinesin population, the FRET efficiency of CLIP-KIF5A-SNAP was more obviously polarised into two populations in the presence of microtubules compared to KIF5A alone (Figure 7A). For KIF5A with microtubule, the majority of FRET bursts have a low FRET efficiency, indicative of an open conformation. There is also a slight increase in the abundance of a high FRET population (FRET efficiency >0.5) indicative of a closed conformation. Concomitant with this change, we observed a depletion of mid-FRET observations that previously created one continuous skewed population of KIF5A FRET efficiency (Figure 3A). The polarisation into two distinct populations was also observable in the cumulative frequency plot (Figure 7B). At lower values of FRET efficiency, the cumulative frequency of KIF5A with microtubules (red) starts to the left of the KIF5A-only population (blue). However, because of the discontinuous distribution of KIF5A FRET in the presence of microtubules, these cumulative frequency plots cross over at higher FRET efficiency values.

**Figure 7:**
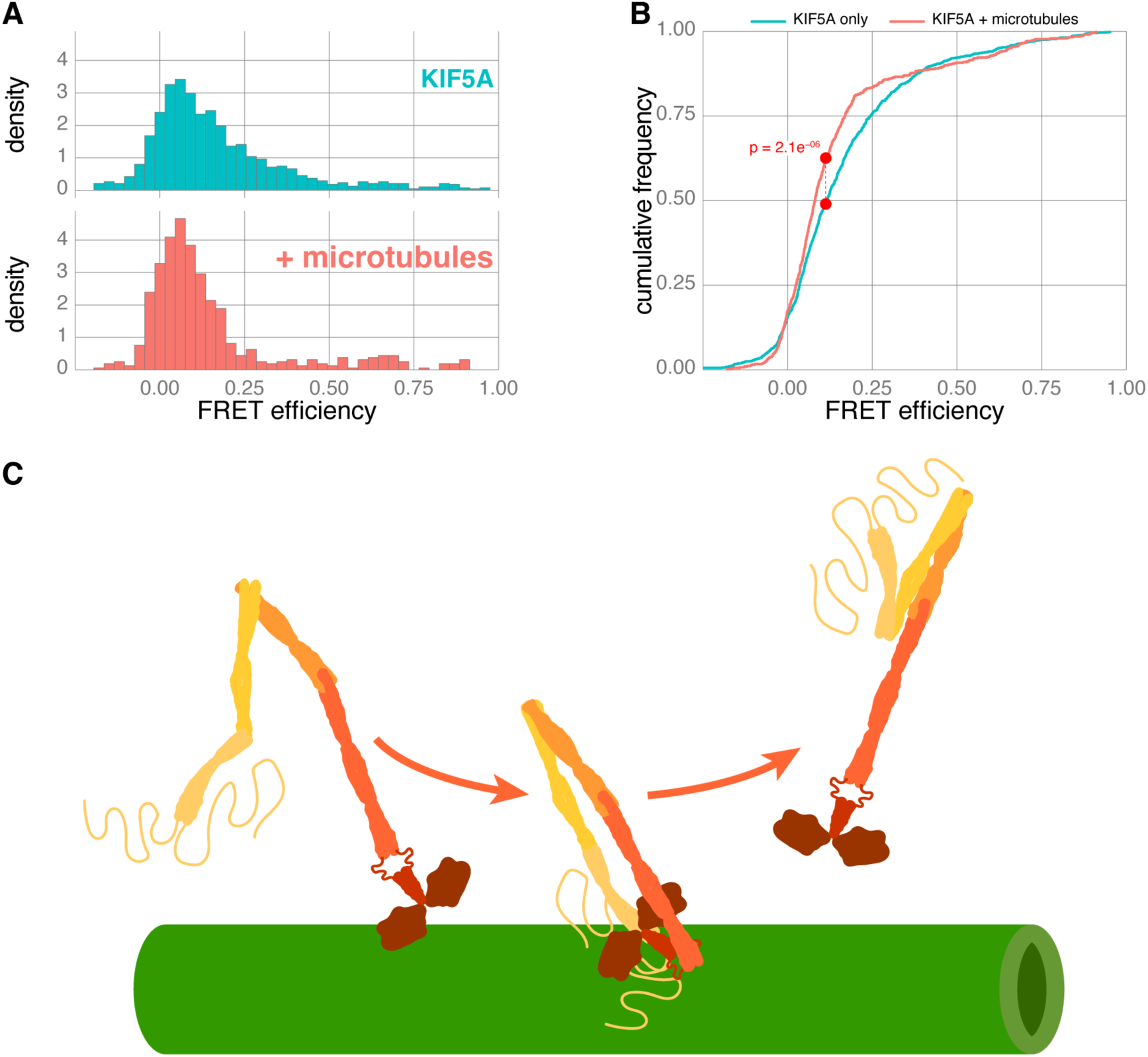
smFRET of KIF5A in the presence of microtubules reveals open and closed conformations. (A) Histograms comparing the FRET efficiencies of KIF5A in the presence (red) and absence (blue) of microtubules. Data combined from 7 biological replicates, n = 529 & 1247 bursts respectively. (B) Cumulative frequency plots comparing FRET efficiencies of KIF5A in the presence and absence of microtubules. (C) Cartoon of kinesin inhibition model in the context of microtubules. Kinesin is autoinhibited in the context of microtubules when not itself bound to adaptors and cargo, but open to binding partners once released from the surface.

Based on our smFRET data in the presence of microtubules, we hypothesise that effective autoinhibition of the kinesin-1 homodimer is far more likely in the context of microtubules (Figure 7C). Indeed the kinesin-1 tail has been known for some time to have microtubule binding motifs, and these could effectively encourage the close apposition of the kinesin-1 head and tail in the context of the microtubule surface and the absence of cargo. Importantly, this model allows us to reconcile both the intact auto inhibition of kinesin-1 observed in TIRFM assays (Figure 2F) and the observation that the majority of in-solution kinesin-1 is in an open conformation (Figures 3 to 6).

## DISCUSSION

We aimed to understand the structural changes that allow the kinesin-1 homodimer to transition between activated and autoinhibited states, by applying biophysical approaches that profile dynamic protein populations rather than static-states. Contrary to expectations of a binary conformational landscape (open vs closed), our findings reveal a broad spectrum of conformations adopted by kinesin-1 homodimers. Ionic strength plays a significant role in modulating this structural landscape. Based on both smFRET and FIDA experiments, we conclude that the majority of kinesin-1 homodimers in solution are in an open, rather than a closed conformation. Interestingly, the presence of microtubules polarised this conformational landscape, potentially facilitating autoinhibition, while simultaneously promoting cargo search. It is important to note that the predominantly open conformation of kinesin-1 homodimers we observed here is not an impediment to effective autoinhibition, as this is still robust in TIRFM microtubule walking assays using similar assay conditions (Figure 2F). Consequently, we demonstrate that solution based measurements can unlock valuable new information about the mechanisms underlying kinesin-1 homodimer function and regulation.

### A versatile expression and labelling platform for single molecule studies

To bridge the gap between *in vitro* and *in vivo* studies of molecular machines, we developed and benchmarked a versatile expression and labelling approach in HEK 293 cells. This approach has not previously been used for smFRET measurements, but we were inspired by the increasing popularity of *in vitro* motility assays using cell lysate as an important validation step in interpreting motor behaviour within complex systems ^18,23,45,48,49^. All samples for smFRET were derived from a single 3.5 cm or 10 cm diameter well/dish using standard tissue culture procedures and commercially available cell permeable fluorescent ligands. This approach enables the production of high-quality protein in sufficient quantities for both single-molecule measurements and downstream analyses like in-gel fluorescence. Using this method, we can easily obtain biological replicates and avoid relying on a single largescale protein purification. Additionally, HEK 293 cells are highly efficient at producing dimeric protein from one purification step with negligible degradation products (as demonstrated by mass photometry, Figure 2H), eliminating the need for further purification steps. This approach was also readily able to produce protein for crosslinking and flash-freezing experiments (Figure 6), as well as labelling with brighter, cell-impermeable, fluorophores (Figure 4E), using one step purification approaches. Beyond costeffectiveness, this method opens up possibilities for studying protein responses to changes in the cellular environment in the future, such as kinase activation or alterations in gene expression. More importantly, we hope that by combining ready expression with an optimised labelling system and increasingly accessible technology platforms ^50^, it will lower the barrier to entry for biophysical characterisation for many proteins and their complexes.

### In solution measurements of kinesin-1 conformational ensembles

The interpretation of our results has been enhanced by combining both ensemble and single molecule in solution measurement techniques. Whereas smFRET accumulates information from observations of individual molecules ∼2 ms in duration, FIDA analysis is based on the observation of a population ensemble ∼2 minutes in duration. By highlighting a decrease in the hydrodynamic radius, FIDA allows us to observe the large overall conformational change of kinesin-1 homodimers when the NaCl concentration is raised from 0 to 150 mM, while parallel smFRET tells us this conformational change likely comes from closer association of the motor and tail of kinesin-1. As it does not average measurements over time or populations - a key advantage of single molecule approaches - smFRET also reveals that even at 150 mM NaCl, kinesin-1 still exists as a conformational ensemble rather than a stable conformation - with the equilibrium in this conformational landscape shifted to make the head-to-tail proximity more likely.

Soon after the discovery of kinesin-1^51,52^, it was observed that changing ionic strength could produce large conformational changes. Hisanaga et al., used low angle rotary shadowing EM and kinesin-1 purified from adrenal glands, to observe conformational diversity and a tendency to change from compact to elongated states in response to increasing ionic strength ^26^. Similar conformational changes were observed in ensemble measurements by sucrose gradient fractionation, using kinesin-1 purified from bovine brain ^27^ and later recombinant drosophila KHC ^34,46^. The major difference between our own work and these original studies is the observation of an additional open conformation at low ionic strength, before closed conformations become more dominant at 150 mM NaCl. These differences are likely due to substantial differences in sample preparation strategies, but also buffer composition and baseline ionic strength. On the other hand we found it difficult to observe kinesin-1 in the presence of 800 mM NaCl by smFRET, likely because of the salting out effect and longer acquisition times needed for smFRET measurements. Salting out could readily be observed by the presence of an increasing number of aggregates by FIDA (Figure 4E).

During this investigation we noticed that kinesin behaviour is extremely sensitive not only to ionic concentration, but also to buffering salts. The sensitivity of kinesin to its environment is also likely behind the very different size exclusion chromatography profiles generated during kinesin-1 purification by different research teams ^17,18,33^. KIF5A in particular seems to have a stronger affinity for self association under certain conditions ^19,33^, however we did not observe this within our own experimental procedures. Although a 150 mM NaCl environment is improbable within the cytosol, kinesin-1’s sensitivity to ionic strength hints at a potential response to physiological ion fluctuations, such as calcium, which was previously thought to be solely regulated by kinesin-interacting proteins like Miro1^53^.

### The changing landscape of kinesin-1 autoinhibition

The head to tail interaction of kinesin-1 was originally linked to its autoinhibition through both single molecule motility ^34,44^ and ATPase assays ^31,46^. This was subsequently validated by structural studies on isolated motor domains and peptides containing the conserved IAK motif from the tail ^29,30^. More recently the dominant role of the IAK motif in preventing motility has been questioned. Mutating the IAK motif alone has little impact on enhancing the motility of kinesin-1 homodimers using *in vitro* motility assays ^18,33^. Similarly, cross-linking mass spectrometry of KIF5B and KIF5C did not reveal contacts between the IAK motif and kinesin motor domains, and the mutation of IAK was not sufficient to prevent the folding of kinesin-1^18^. However, a similar cross-linking mass spectrometry study of KIF5A did find interactions between the motor domains and the IAK motif ^19^ and recent cryo-EM work with chimeric KIF5B molecules (consisting of only motor domains and IAK containing tail), does show the IAK docked between the motor domains when bound to microtubules ^32^. Given the large scale conformational changes we have been able to observe here in the absence of microtubules, it seems highly unlikely that the IAK motif is solely responsible for opening and closing the kinesin motor, but it is still likely to have a direct role in the molecular mechanism of autoinhibition in response to microtubules.

AlphaFold predicts an interlocking of CC1 and CC2 to form a retrograde loop in the location previously ascribed to be a hinge in kinesin-1^34,44^. In our experience this prediction has been consistent across both AlphaFold versions and species (e.g. D. *melanogaster*, C. *elegans, Loligo pealeii* and *Strongylocentrotus purpuratus*). Three recent low resolution negative stain EM studies captured the closed conformation of different isoforms of the full length kinesin-1 motor ^17–19^. However, while providing valuable information, all studies used crosslinking to enrich the closed state and lack the high resolution structural information to confirm the AlphaFold prediction of a more rigid (rather than hinge-like) CC1-CC2 region. *In vitro* motility and ATPase assays provided some of the original evidence that the region between CC1 and CC2 was the main region of flexibility mediating the head to tail interaction of kinesin-1^34,44^. However, more recent results show a very limited impact on increasing motility when deleting this region, compared to constitutively active controls ^33,45^. Our data suggests that the Δ505-610 mutant is still highly flexible and likely capable of making head-to-tail interactions, having a FRET efficiency profile highly similar to full length KIF5A. This is supported by previous results on a similar hinge mutant that formed a compact structure by the comparison of sedimentation coefficients ^34^. The overall flexibility of kinesin-1 along its length likely underpins the limited impact of mutating any one flexible region, hinge 1 or hinge 2 (Figure 2B), observed using *in vitro* motility assays ^18^. In support of a more rigid protein structure interlocking CC1 and CC2, we observed the lowest FRET efficiency in our study for the KIF5A(411-680) construct, a CC1-CC2 fragment (Figure 6D & Figure 6H). If residues 564-580 were a true hinge allowing large conformational changes in this region, we would expect a larger proportion of events to occupy higher FRET efficiency values. Consequently we conclude that AlphaFold predictions are likely correct, however it will be interesting to test if the interlocking coiled coils of CC1-CC2 can be separated by mutation or post translational modification in the future.

Although we did not not study the impact of KLCs on the conformational landscape of kinesin-1 in this study, the concertina structure currently predicted by AlphaFold and presented here supports the role of KLCs in promoting a closed conformation of kinesin-1. The KLC binding site spans CC3 and CC4, and is thought to stabilise the region of flexibility (hinge 2, Figure 2B) in between ^18^. It has been long established that the basal ATPase rate of kinesins with KLCs bound is lower than the heavy chains alone ^24,34^. Supporting evidence also points to a role of KLCs in favouring an autoinhibited conformation, in cells ^39,54^, through crosslinking studies ^18^ and *in vitro* motility assays ^33^. However, both open and closed states of kinesin-1 heterotetramers (kinesin-1 with additional light chains bound) can still be separated by size exclusion chromatography ^17,18^, so it will be important to understand how the presence of KLCs alters the conformational landscape of kinesin-1.

Our work suggests that kinesin-1 does not have to be in a permanently closed conformation to have an effective autoinhibition mechanism. Kinesin is fundamentally a microtubule-stimulated ATPase ^28^; if the time spent not bound to cargo and microtubules is insignificant, then ATPase activity away from the microtubule surface may be sufficiently low so as to not require further inhibition in the cellular context. From many *in vitro* studies to date, it is also clear that binding to microtubules is severely inhibited when kinesin isn’t bound to cargo ^21,22,33^. Consequently it’s already safe to assume that cells tolerate the basel level of ATPase activity that kinesin is capable of when it is bound to cargo, but yet to find a microtubule. The other implication of our results is that binding to microtubules promotes the autoinhibited conformation when cargo is absent. Previous work proposed that the interaction of the kinesin tail with microtubules could support kinesin ‘pausing’ ^29^, whereas it could have a more active role in autoinhibition. It will be interesting to investigate whether the previously identified ATP independent microtubule binding site preceding the IAK motif has a role in this inhibition ^31,54–56^.

### The cellular role for dynamic structural landscape of kinesin-1

Kinesin-1 was discovered due to its essential role in fast axonal transport ^51,52^. In the intervening years it has also become established that many modes of slow axonal transport are also dependent on kinesin-1^10,11,57^. Whereas fast axonal transport proceeds at close to the maximum speed of processive kinesins ^58^, slow transport is ∼10 to ∼100 fold slower ^59^. There is currently no biochemical or biophysical explanation for kinesin-1 activation by cargo that explains this discrepancy. We recently proposed that one possible mechanism for slow axonal transport is a rate limiting supply of kinesin for certain classes of cargo ^10^. This allows for short bursts of motility in a cellular environment where many low affinity cargos compete for a few free motors. However, if kinesin primarily exists in a stable locked down state, it is difficult to envision how this could be the case; the same series of high affinity interactions would be needed to unlock the motor, regardless of whether a cargo needed to stably recruit a motor, or only hang on for a short burst. Due to the lack of high resolution structural data, structural conformations of kinesin have often been inferred from *in vitro* and in cell activity assays. This could have masked the intrinsic conformational heterogeneity of kinesin-1, but the dynamics of the conformational ensembles are likely to be essential to kinesin-1 function in cells. A conformational ensemble, where kinesin-1 is constantly shifting between open and closed states, allows for much greater diversity of cargo interactions; both stable recruitment and a more transient but promiscuous mode depending on the cellular environment.

Our hypothesis moving forward is that cargo recognition is about stabilising the open conformation, rather than unfolding and ‘activating’ kinesin, at least where KLC independent functions of kinesin-1 are concerned. Further, that preventing a head-to-tail interaction would be sufficient to promote a basal level of active transport. With this in mind, more possible scenarios of cargo recruitment are possible that could be better suited to the diverse array of cellular transport functions attributed to kinesin-1. For example, AlphaFold has been consistent in its predictions of two key regions of flexibility, hinge 1 and 2. This raises the possibility that the activity of kinesin-1 could be promoted through either (i) stabilising hinge 1 and 2 into a fully elongated structure, or (ii) stabilising the interaction of CC4 with CC3. In support of the first model, we previously predicted stabilisation of an additional point of flexibility in the kinesin-1 homodimer, based on the activation of KIF5C by cooperation between the adaptors HAP1 and GRIP1^21^. We believe that by accounting for the intrinsic dynamics of kinesin dimers, solution based measurements to study kinesin-1 conformational ensembles will be essential to building a comprehensive picture of both autoinhibition and cargo binding.

## REAGENTS AND TOOLS TABLE

Key reagents and their sources are listed in Table 1.

**Table 1:** Reagents and tools Key reagents used in this study and their source.

| Reagent/Resource | Reference or Source | Identifier or Catalog Number |
| --- | --- | --- |
| **Experimental models** |  |  |
| HEK293 cells line | ECACC | 85120602 |
| 293FT cell line | ThermoFisher Scientific | R70007 |
| **Recombinant DNA** |  |  |
| pCLAP (pCDNA3.1-CLIP-SNAP-HisTag-FLAG) | this study | Addgene #200786 |
| pCDNA3.1-CLIP | this study | Addgene #200787 |
| pCDNA3.1-SNAP | this study | Addgene #200788 |
| pCDN3.1-StrepTag-HaloTag | this study | Addgene #219682 |
| CLIP-KIF5A-SNAP-HisTag-FLAG (pCLAP-KIF5A) | this study | Addgene #200789 |
| pCMV-KIF5B(1-560)-HaloTag-StrepTag (K560-Halo) | this study | Addgene #219681 |
| pCDNA3.1-CLIP-KIF5A(1-573) | this study | Addgene #219683 |
| pCDNA3.1-CLIP-KIF5A | this study | Addgene #200790 |
| pCDNA3.1-KIF5A-SNAP-HisTag-FLAG | this study | Addgene #200791 |
| pCDNA3.1-CLIP-KIF5A(954-1027)-SNAP-HisTag-FLAG (pCLAP-KIF5A(954-1027)) | this study | Addgene #200792 |
| pCDNA3.1-CLIP-KIF5A(1-915;505-610)-SNAP-HisTag-FLAG (pCLAP-KIF5A(1-915;505-610)) | this study | Addgene #200793 |
| pCDNA3.1-CLIP-KIF5A(411-680)-SNAP-HisTag-FLAG (pCLAP-KIF5A(411-680)) | this study | Addgene #200798 |
| pCDNA3.1-CLIP-KIF5B-SNAP-HisTag-FLAG (pCLAP-KIF5B) | this study | Addgene #200799 |
| **Antibodies** |  |  |
| Alexa Fluor 680-AffiniPure Donkey Anti-Rabbit IgG (H+L) | Jackson ImmunoResearch Laboratories, Inc. | 711-625-152 |
| Alexa Fluor 790-AffiniPure Donkey Anti-Mouse IgG (H+L) | Jackson ImmunoResearch Laboratories, Inc. | 715-655-150 |
| Mouse anti-GAPDH | Proteintech | 60004-1-1g |
| Mouse anti-β-Tubulin antibody | Sigma-Aldrich | T5201-100UL |
| Rabbit anti-DYKDDDDK | Proteintech | 20543-1-AP |
| Rabbit anti-KIF5B | Proteintech | 21632-1-AP |
| **Chemicals, Enzymes and other reagents** |  |  |
| 3xDYKDDDDK-peptide | Proteintech | fp-1 |
| BS3 (bis(sulfosuccinimidyl)suberate) | ThermoFisher Scientific | 16091992 |
| BSA | Sigma-Aldrich | AO281 |
| CLIP-Cell TMR-Star | New England Biolabs | S9219S |
| DYKDDDDK Fab-Trap Agarose | Proteintech | ffa-20 |
| Dynabead+ProtG | ThermoFisher Scientific | 10003D |
| Fluorescent tubulin (HiLyte488) | Cytoskeleton | TL488M |
| HaloTag TMR Ligand | Promega | G8251 |
| Paclitaxel | Cambridge Bioscience | CAY10461 |
| Porcine brain tubulin | Cytoskeleton | T240 |
| SNAP-Cell 647-SiR | New England Biolabs | S9102S |
| SNAP-Surface Alexa Fluor 488 | New England Biolabs | S9129S |
| SNAP-Cell TMR-Star | New England Biolabs | S9105S |
| LipoD293 DNA In Vitro Transfection Reagent | SignaGen Laboratories | SL100668 |
| Pluronic F-127 | Sigma-Aldrich | P2443-250G |
| Adenosine 5-triphosphate (ATP) disodium salt hydrate | Sigma-Aldrich | A2383-1G |
| Glucose oxidase | Scientific Laboratory Supplies | G2133-50KU |
| Catalase | Sigma-Aldrich | C3155-50MG |
| **Software** |  |  |
| Jupyter Notebook | https://jupyter.org/ |  |
| FRETbursts python package | https://fretbursts.readthedocs.io/en/latest/ |  |
| RStudio | https://cran.r-project.org/ |  |
| DiscoverMP | https://www.refeyn.com/ |  |
| Fida software 3.0 | https://www.fidabio.com/ |  |
| Fiji/ImageJ | https://imagej.net/software/fiji/ |  |
| TrackMate v7.11.1 | 64 |  |
| AlphaFold v2.3.2 | 67 |  |
| **Other** |  |  |
| BD FACS Melody | BD Biosciences |  |
| Odyssey Fc Imager | LI-COR |  |
| Custom built smfBox | 50 |  |
| EI-FLEX | Exciting Instruments |  |
| TwoMP mass photometer | Refeyn |  |
| Fida Neo | FidaBio |  |
| Ti-Ns N-STORM | Nikon |  |
| Custom built single-molecule scanning TIRF/FRAP imaging system | Cairn research |  |

## MATERIALS AND METHODS

### Recombinant DNA expression plasmids

All plasmids used in the smFRET assays were based on a pCDNA3.1-CLIP-SNAP-HisTag-FLAG plasmid we designed and named pCLAP. The CLIP tag is derived from the SNAP tag, with only eight amino acid changes, meaning DNA sequences are highly similar. To use both tags in the same construct, we codon optimised CLIP and SNAP linked by the short flexible linker sequence G-S-A-G-S-A-A-G-S-G-E-F ^60^ for human expression, such that stretches of identity between the two nucleotide sequences were no longer than nine base pairs. Kinesin-1 (mouse KIF5A or human KIF5B) was inserted between the CLIP and SNAP tags creating CLIP-KIF5A-SNAP and CLIP-KIF5B-SNAP. Kinesin truncation mutants or single tagged constructs were generated by deletion mutagenesis PCR from these plasmids. Similarly, CLIP and SNAP only plasmids were generated by deletion mutagenesis PCR of pCLAP. All constructs with the SNAP tag retain the C-terminal His and FLAG tags for protein purification and immuno-assays. The constitutively active human KIF5B construct (K560), C-terminally tagged with Halo and Strep tags, was derived from the previously published K560-HaloTag ^10^ through inframe insertion of the Strep-tag at the C-terminus. The StrepTag-HaloTag construct was codon optimised for human expression and is used to show non-specific labelling of the HaloTag (Figure S1H). All plasmids are available from Addgene (see Materials availability).

### Cell culture and transfection

HEK 293 (Human Embryo Kidney 293) cells, were cultured in Dulbecco’s Modified Eagle Medium (DMEM) supplemented with 10 % (vol/vol) Fetal Bovine Serum (FBS), 500 U Penicillin and 500 *µ*g Streptomycin. 293FT cells (a clonal derivative of the Human Embryo Kidney 293 cell line), were cultured in DMEM supplemented with 10 % FBS, 2 mM GlutaMAX, 500 *µ*g/ml Geneticin, 1 mM Sodium Pyruvate, 1 X MEM Non-Essential Amino Acid Solution, 500 U Penicillin and 500 *µ*g Streptomycin. Both cell lines were grown at 37 °C with 5 % CO_2_ and passaged every 2-3 days when ∼80% confluent by trypsinization and replating. Both cell lines were checked regularly for mycoplasma contamination.

18 hours after plating at ∼50 % confluency in either a 6 well plate or 10 cm dish, cells were transfected using the LipoD293 DNA In Vitro Transfection Reagent. For a 6 well plate, media was removed so that only 1 ml was covering the cells. A volume of DNA equal to 1 *µ*g was gently mixed into 50 *µ*l of DMEM whilst 3 *µ*l of transfection reagent was mixed into 50 *µ*l of DMEM before being added dropwise to the diluted DNA solution. The DNA mixture was left to incubate at room temperature for 10 minutes before adding to the well in a dropwise manner. 24 hours after transfection, cells were washed with 1 ml Phosphate Buffered Saline (PBS) and lifted with 500 *µ*l of 0.5 % trypsin for ∼ 5 minutes. 1 ml of media was added to neutralise the trypsin and cells were harvested by centrifugation at 500 x g for 4 minutes at 4 °C. The cell pellet was resuspended in 1 ml PBS before a further centrifugation at 500 x g at 4 °C. All volumes were increased by a factor of five for a 10 cm dish.

### Flow cytometry analysis of ligand washout conditions

Untransfected HEK 293 cells were plated at ∼50 % confluency into a six well plate. 24 hours after plating, CLIP-Cell TMR-Star or SNAP-Cell 647-SiR was added to the cells at a final concentration of 0.25 *µ*M and incubated for one hour. The cells were washed three times with media and either harvested immediately or left to incubate for a further 1 to 4 hours. Cells were harvested (as described above) and the pellet resuspended in 0.5 ml PBS. The cell suspension was analysed using a BD FACS Melody flow cytometer (CLIP-Cell TMR-Star - 561 nm laser, PE 582/15 filter or SNAP-Cell 647-SiR - 633 nm laser, APC 660/10 filter). The raw data was transformed and the fluorescence intensity plotted using a custom R notebook (see data and code availability) and the flowcore package 61. The data from each biological repeat was combined and the median value for each sample compared.

### In-gel fluorescence analysis of ligand labelling

HEK 293 cells were plated into 6-well plates and transfected with either CLIP or SNAP (as described above). To optimise the dye incubation time, 24 hours after transfection CLIP-Cell TMR-Star or SNAP-Cell 647-SiR was added to the cells for five different incubation periods (15 minutes, 30 minutes, 1 hour, 2 hours and overnight) at a final concentration of 0.25 *µ*M. To optimise the dye concentration, 24 hours after transfection CLIP-Cell TMR-Star or SNAP-Cell 647-SiR was added to cells at five different concentrations (0.1 *µ*M, 0.25 *µ*M, 0.5 *µ*M, 1 *µ*M and 2 *µ*M) and left to incubate according to the previously optimised incubation conditions (overnight for CLIP-Cell TMR-Star and 2 hours for SNAP-Cell 647-SiR). To determine the cross-reactivity between the two dyes, 24 hours after transfection 0.25 *µ*M of CLIP-Cell TMR-Star and SNAP-Cell 647-SiR were added to the cells and left to incubate either for 2 hours or overnight. For all these experiments, cells were harvested (as described above) and lysed in 100 *µ*l of lysis buffer (40 mM HEPES pH 7.5, 1 mM EDTA pH 8, 120 mM NaCl, 0.05 % Triton X-100, 1 *µ*g/ml Aprotinin, 10 *µ*g/ml Leupeptin, 1 *µ*g/ml Pepstatin A and 10 *µ*g/ml tosyl-L-Arginine Methyl Ester) before incubation on ice for 10 minutes. Lysates were clarified by centrifugation at 17,000 x g for 10 minutes at 4 °C. 3X sample buffer (150 mM Tris pH 6.8, 6% SDS, 0.3 M DTT, 0.3 % Bromophenol and 30% glycerol) was added to the resultant supernatant to a final 1X concentration. 20 *µ*l of each sample was resolved by SDS-PAGE on 15 % polyacrylamide resolving gels at 150V for ∼ two hours. In-gel fluorescence of bound ligands was captured using the Odyssey Fc Imager (LI-COR) in the 600 nm and 700 nm channels. The signal intensity for each band was normalised to the lowest incubation or concentration per biological repeat and plotted using R (see Data and code availability, below).

### Immunoprecipitation and western blotting

HEK 293 cells were plated into 6-well plates and transfected as described above. 24 hours after transfection, media was removed and the cells scraped into 1.5 ml PBS. Cells were pelleted by centrifugation at 400 x g for two minutes at room temperature. The cell pellet was washed by resuspening in 1 ml PBS and centrifugation at 2300 x g for five minutes at 4 °C. Cells were lysed by resuspending the pellet in 300 *µ*l of lysis buffer (50 mM HEPES pH 7.5, 1 mM EDTA pH 8, 1 mM MgCl_2_, 25 mM NaCl, 0.5 % Triton-X-100, 1 mM DTT, 1 *µ*g/ml Aprotinin, 10 *µ*g/ml Leupeptin, 1 *µ*g/ml Pepstatin A and 10 *µ*g/ml tosyl-L-Arginine Methyl Ester) and incubating on ice for 10 minutes. Lysates were clarified by centrifugation at 17,000 x g for 10 minutes at 4 °C. 1 *µ*g of rabbit anti-DYKDDDDK was added to the supernatant and incubated with rotation for an hour at 4 °C. 25 *µ*l of Dynabead+ProtG beads were washed three times by addition of 200 *µ*l of PBS + 0.02 % Tween and discarding the supernatant. The cell lysate antibody mix was added to the beads and incubated with rotation for 10 minutes at room temperature. The supernatant was removed and the beads washed three times with 200 *µ*l of PBS before resuspending in 1X sample buffer (50 mM Tris pH 6.8, 2% SDS, 0.1 M DTT, 0.1 % Bromophenol and 10% glycerol). 20 *µ*l of each sample was resolved by SDS-PAGE on a 10 % polyacrylamide resolving gel and transferred to Immobilon-FL PVDF membrane. Membranes were probed with primary antibodies rabbit anti-DYKDDDDK, rabbit anti-KIF5B or mouse anti-GAPDH as indicated; fluorescent secondaries were Alexa Fluor 680-AffiniPure Donkey Anti-Rabbit or Alexa Fluor 790-AffiniPure Donkey Anti-Mouse. Fluorescent western membranes were visualised on the Odyssey Fc Imager (LI-COR).

### smFRET assays

Six hours after transfection, CLIP-Cell TMR-Star was added to the HEK 293 cells in 6-well plates at a final concentration of 0.5 *µ*M and left to incubate for around 15 hours. SNAP-Cell 647-SiR (added at the same final concentration) was added to the cells for a further two hours. The cells were then washed three times with media and incubated in 3 ml of fresh media for two hours to remove any unbound dye. Cells were harvested as above and lysed in 50 *µ*l of lysis buffer (40 mM HEPES pH 7.5, 1 mM EDTA pH 8, 120 mM NaCl, 0.05 % Triton-X-100, 0.1 mg/ml photobleached BSA, 1 mM DTT, 1 mM Mg.ATP, 1 *µ*g/ml Aprotinin, 10 *µ*g/ml Leupeptin, 1 *µ*g/ml Pepstatin A and 10 *µ*g/ml tosyl-L-Arginine Methyl Ester) before incubation on ice for 10 minutes. Lysates were clarified by centrifugation at 17,000 x g for 10 minutes at 4 °C. The supernatant was diluted in smFRET assay buffer (12 mM PIPES pH 6.7, 1 mM EGTA pH 7, 2 mM MgCl_2_, 0.1 mg/ml photobleached BSA, 1 mM DTT and 1 mM Mg.ATP) to result in single molecule resolution (usually 1:100,000). The sample was applied either to a custom built smFRET confocal microscope (smfBox) described in detail previously 50, or an EI-FLEX with the same configuration. Briefly, the smfBox consists of two lasers (80 mW 515 nm and 100 mW 638 nm), two emission filters (571/72 and 678/41) and two avalanche photodiodes; for the donor and acceptor respectively. Data collection was typically in three 15 minute periods per sample. Data was analysed using a custom built Jupyter (for initial elimination of background and burst selection using dye specific correction factors using the FRETBursts package 62) and R (for burst filtration and plotting) notebooks (see Data and code availability, below). The burst threshold was set at 20 photons in both the DD+DA and AA channels for the burst selection. Any technical repeats with a burst rate above 1 burst per second were eliminated and the background filters were set as above: 8.5 photons for donor excitation (DD+DA), 2.6 for donor emission (DD) and 5 for acceptor excitation and emission (AA). Displayed p values for paired comparisons were calculated by Kolmogorov-Smirnov test and are shown plotted at the point of maximum difference between samples. For multiple comparisons, pairwise Kolmogorov-Smirnov tests were performed with Bonferroni correction. All labelled samples used for smFRET were also checked using in-gel fluorescence of samples resolved by SDS-PAGE on a 10 % polyacrylamide gel followed by visualisation on an Odyssey Fc Imager (LI-COR).

### Small batch protein purification and cross-linking

24 hours after transfection, 10 cm dishes of 293FT cells were harvested by removing the media and the cells scraped into 5 ml PBS. Cells were pelleted by centrifugation at 400 x g for two minutes at room temperature and washed by resuspening in 1 ml PBS. After centrifugation at 2300 x g for five minutes at 4 °C the resultant pellet was resuspended in 200 *µ*l lysis buffer (50 mM HEPES pH 7.5, 1 mM EDTA pH 8, 1 mM MgCl_2_, 25 mM KAc, 0.5 % Triton-X-100, 1 mM DTT, 1 *µ*g/ml Aprotinin, 10 *µ*g/ml Leupeptin, 1 *µ*g/ml Pepstatin A and 10 *µ*g/ml tosyl-L-Arginine Methyl Ester). 25 *µ*l of DYKDDDDK Fab-Trap Agarose beads were washed three times by addition of 500 *µ*l dilution buffer (50 mM HEPES pH 7.5, 1 mM EGTA pH 7, 2 mM MgCl_2_ and 150 mM KAc), pelleting by centrifugation at 2500 x g for 5 minutes at 4 °C and discarding the supernatant. The lysate (diluted in 300 *µ*l dilution buffer) was equilibrated with the washed beads by incubation with rotation for one hour at 4 °C. For on bead labelling, 2 *µ*l of each ligand (CLIP-Cell TMR-Star and SNAP-Cell 647-SiR, 600 *µ*M stock) was also added to the lysate. After incubation, the beads were washed three times by centrifugation at 2500 x g for 5 minutes at 4 °C and resuspension in 500 *µ*l wash buffer (50 mM HEPES pH 7.5, 1 mM EDTA pH 8, 2 mM MgCl_2_, 150 mM KAc, 1 mM DTT, 1 *µ*g/ml Aprotinin, 10 *µ*g/ml Leupeptin, 1 *µ*g/ml Pepstatin A and 10 *µ*g/ml tosyl-L-Arginine Methyl Ester). 3xDYKDDDDK-peptide was diluted to 150 *µ*g/ml with PBS and 100 *µ*l added to the washed beads. This suspension was incubated with rotation for 20 minutes at room temperature. Eluted protein was separated from beads by centrifugation at 2500 x g for 2 minutes at 4 °C. Protein concentration was determined by BCA assay. If cross-linking was required, BS3 was diluted to 2 mg/ml in the dilution buffer and added to half the sample at a 1:1 mass ratio. The mixture was incubated on ice for twenty minutes, before cross-linking was quenched by addition of Tris pH 7.5 to a final concentration of 25 mM and further incubated on ice for twenty minutes. Gel samples of each stage were analysed by in-gel fluorescence with 20 *µ*l of each sample resolved by SDS-PAGE with a 10 % polyacrylamide resolving gel. Following fluorescence imaging, the gel was stained with Quick Coomassie Stain (Protein Ark) and total protein imaged for comparison.

### Mass photometry

Samples were prepared as above by small batch protein purification without addition of the crosslinker and analysed using the Refeyn TwoMP mass photometer. Measurements were carried out at room temperature. All resultant images were analysed using the Refeyn DiscoverMP software and conversion from interferometric contrast to molecular weights was calibrated using Refeyn’s protein standards. Gaussian fitting to mass histograms was performed using R (see Data and code availability, below).

### Flow induced dispersion analysis

Samples were prepared as above by small batch protein purification without addition of the crosslinker. To label the protein, 2 *µ*l of 0.6 mM SNAP-Surface Alexa Fluor 488 was added to the lysate before addition to the beads. Protein was analysed using a Fida Neo instrument with an excitation wavelength of 480 nm. Purified CLIP-KIF5A-SNAP was mixed with the corresponding BRB80 buffer (80 mM PIPES pH 6.7, 1 mM EGTA pH 7, 2 mM MgCl_2_) supplemented with the relevant NaCl concentration (0 - 1 M) within the instrument, immediately prior to measurement. Each salt concentration was run in triplicate. The resultant Taylorgrams were fitted using the standard parameters and smooth curve settings in the FIDA software (version 3.0) to determine hydrodynamic radii and spike count for each condition. This data was plotted using R (see Data and code availability, below).

### Microtubule polymerisation for *in vitro* assays

Lyophilised tubulins (both unlabelled and HiLyte 488 labelled) were resuspended to 5 mg/ml in GBRB80 (1 mM GTP, 80 mM PIPES pH 6.7, 1 mM EGTA pH 7, 2 mM MgCl_2_), flash frozen and stored at -80 °C. 40 *µ*l of this tubulin stock (0.8:40 ratio of labelled:unlabelled tubulin if making fluorescent microtubules) was thawed on ice and centrifuged at 270,000 g (TLA-100.3 rotor) for 10 minutes at 4 °C. The supernatant was then incubated at 37 °C for twenty minutes before addition of Paclitaxel to a final concentration of 40 *µ*M and further incubation for twenty minutes. Microtubules were pelleted by centrifugation at 270,000 x g (TLA-100.3 rotor) for 10 minutes at 4 °C and gently resuspended in TBRB80 (40 *µ*M Paclitaxel, 80 mM PIPES pH 6.7, 1 mM EGTA pH 7, 2 mM MgCl_2_). Microtubules were kept in the dark at room temperature until use. Microtubules used in smFRET assays were sheared by passing through a Hamilton 25 *µ*l 22G fixed needle syringe ten times before diluting 1 *µ*l in 250 *µ*l smFRET assay buffer (12 mM PIPES pH 6.7, 1 mM EGTA pH 7, 2 mM MgCl_2_, 0.1 mg/ml photobleached BSA, 1 mM DTT and 1 mM Mg.ATP) and using this to dilute the CLIP-KIF5A-SNAP transfected cell lysate.

### Single molecule motility assays

24 hours after transfection, HaloTag TMR or SNAP-Cell TMR-Star ligands were added to 293FT cells at a final concentration of 25 nM and 0.5 *µ*M respectively. Cells labelled with HaloTag TMR were incubated for 15 minutes and those with SNAP-Cell TMR-Star for 2 hours. The cells were then washed three times with fresh media and incubated for a further 30 minutes. Cells were harvested and washed in PBS as stated above for immunoprecipitation. The cell pellet was resuspended in 300 *µ*l of lysis buffer (50 mM Tris pH 8, 25 mM KAc, 1 mM MgCl_2_, 1 mM EDTA, 0.5 % Triton-X-100, 1 mM DTT, 1 *µ*g/ml Aprotinin, 10 *µ*g/ml Leupeptin, 1 *µ*g/ml Pepstatin A and 10 *µ*g/ml tosyl-L-Arginine Methyl Ester) and incubated on ice for 10 minutes before centrifugation at 16,000 x g for 10 minutes at 4 °C. The supernatant was collected and stored on ice for use in flow chambers (below).

Flow chambers were prepared as described previously 21. Briefly, a #1.5 22 x 32 mm cleaned and silanised coverslip was attached to a glass microscope slide using double-sided tape. Chambers were defined using vacuum grease. Flow cells were incubated for five minutes with i) 1 chamber volume (CV) of 20 *µ*g/ml anti-*β*-Tubulin antibody (diluted in BRB80), ii) 1 CV of 50 mg/ml F-127, iii) 2 CV of microtubules diluted in TBRB80 (1:250). Unbound microtubules were removed with a 2 CV BRB80 wash step before adding the final reaction mix of 1 CV formed of 1 *µ*l cell lysate in 20 *µ*l of assay buffer (80 mM PIPES pH 6.7, 1 mM EGTA pH 7, 1 mM MgCl_2_, 0.3 mg/ml casein, 0.3 mg/ml BSA 10 mM DTT, 10 mM ATP, 15 mg/ml glucose, 0.5 *µ*g/ml glucose oxidase and 470U/ml catalase).

Motility assays containing CLIPf-KIF5A-SNAPf (Figure 2F) were imaged using the Nikon Ti-NS N-STORM microscope at the Wolfson Light Microscopy Facility with the SR Apo TIRF 100X objective lens and the Andor iXion ultra EM-CCD camera, with 488 nm and 561 nm lasers. Movies were acquired for 1 minute at 40 ms exposure. KIF5B(1-560)-HaloTag and CLIPfKIF5A(1-573) constructs (Figure 2D) were imaged on a custom single-molecule scanning TIRF/FRAP imaging system built by Cairn Research. This setup uses a Ti2-E automated inverted microscope base (Nikon), ASI automated XY & piezo-Z stage controller, iLas 2 scanning TIRF/FRAP unit (Gataca Systems) with dual-collimation optics, Cairn multi-line laserbank, and a Photometrics Prime 95B sCMOS camera. TIRF was performed with a 488 nm and 561 nm laser, a TIRF quad-band filter cube (TRF89901-EMv2, Chroma) and additional emission clean up provided by a Cairn Optospin filter wheel with ET525/50m or ET595/44m filters for 488 and 561 laser lines respectively. Images were taken in MetaMorph acquisition software using a Nikon CFI Apochromat TIRF 100XC oil (N.A. 1.49, W.D. 0.12 mm) objective lens. Exposure time was 40 ms (25 frames per second) and movies were acquired for 1000 frames.

Motility in image stacks was analysed using Fiji 63 and the TrackMate (v7.11.1) plugin 64. Briefly, individual microtubules were isolated from the field of view using the line selection tool followed by the Straighten tool in Fiji. Kinesin molecules were tracked by running TrackMate on the 561 channel, using the LoG detector, an estimated object diameter of 8 pixels and the LAP tracker function. The detected spots were additionally filtered by their Y position (i.e. proximity to the straightened microtubule). Further downstream analysis was performed in R (see Data and code availability, below).

### Protein Structure Prediction

To predict three-dimensional structures of the target proteins, we used AlphaFold v2.3.2 (unless otherwise stated) 65,66, using the AlphaFold Colab notebook hosted on Google Colaboratory 67. For all structures the relaxation stage was enabled and multimer model recycling was set to a limit of six. The highest scoring prediction was used. Structures and associated files are available from ORDA (see Data and code availability, below).

## Resource availability

### Lead contact

All outputs are available as indicated. Further information and requests for resources and reagents should be directed to and will be fulfilled by the lead contact, Alison Twelvetrees.

### Materials availability

Plasmids generated in this study have been deposited and are available from Addgene (www.addgene.org/Alison_Twelvetrees).

### Data and code availability

All data and code that was used to make figures is deposited in ORDA, the University of Sheffield data repository, powered by Figshare (doi.org/10.15131/shef.data.c.7555284). Example Jupyter and R notebooks used for processing raw smFRET data are also available from ORDA.

## ACKNOWLEDGMENTS

We thank Dominic Bingham, Annie Savage and Rebecca Mighell for generating some of the plasmids used in this study. We also thank Benjamin Ambrose and Elliot M. Steele for support conducting smFRET experiments. We are particularly indebted to Daniel Bose for constant discussion of the concepts underpinning the work and constructive comments on the manuscript. We would also like to thank the teams at FIDABio and Refeyn for helping collect and analyse the FIDA and mass photometry data in this work, as well as the Wolfson Light Microscopy Facility and the Flow Cytometry Facility at the University of Sheffield.

This research was supported by the following funding: A.E.T. is a Sir Henry Dale Fellow, funded by the Wellcome Trust and the Royal Society (grant number: 220192/Z/20/Z); E.R.S. was supported by a studentship from the White Rose BBSRC Doctoral Training Partnership (grant number: 2109768); E.T. was supported by a University of Sheffield PGT to PGR Scholarship from the Faculty of Health. E.R.S. also received a University of Sheffield Postgraduate Research Student Publication Scholarship. The N-STORM microscope in the Wolfson Light Microscopy Facility was funded by MRC grant MK/K0157531/1.

## AUTHOR CONTRIBUTIONS

Conceptualization: A.E.T., T.D.C., E.R.S.; Data curation: E.R.S., A.E.T.; Formal analysis: E.R.S., A.E.T.; Funding acquisition: A.E.T., T.D.C., E.R.S.; Investigation: E.R.S., E.T.; Methodology: A.E.T, T.D.C., E.R.S.; Project administration: A.E.T; Supervision: A.E.T, T.D.C.; Resources: A.E.T, T.D.C.; Validation: E.R.S., A.E.T., M.A.S.A; Visualisation: A.E.T.; Writing - original draft: A.E.T.; Writing - review and editing: all authors

## REFERENCES

[1] Cláudia Rosa-Ferreira and Sean Munro. Arl8 and SKIP act together to link lysosomes to kinesin-1. Dev. Cell, 21(6):1171–1178, December 2011.

[2] Elizabeth E Glater, Laura J Megeath, R Steven Stowers, and Thomas L Schwarz. Axonal transport of mitochondria requires milton to recruit kinesin heavy chain and is light chain independent. J. Cell Biol., 173(4):545–557, May 2006.

[3] Marcin J Woźniak, Becky Bola, Kim Brownhill, Yen-Ching Yang, Vesselina Levakova, and Victoria J Allan. Role of kinesin-1 and cytoplasmic dynein in endoplasmic reticulum movement in VERO cells. J. Cell Sci., 122(Pt 12):1979–1989, June 2009.

[4] Alison Emma Twelvetrees, Eunice Y Yuen, I Lorena Arancibia-Carcamo, Andrew F Macaskill, Philippe P Rostaing, Michael J Lumb, Sandrine S Humbert, Antoine A Triller, Frédéric Saudou, Zhen Z Yan, and Josef T Kittler. Delivery of GABAARs to synapses is mediated by HAP1-KIF5 and disrupted by mutant huntingtin. Neuron, 65(1):53–65, January 2010.

[5] Comert Kural, Hwajin Kim, Sheyum Syed, Gohta Goshima, Vladimir I Gelfand, and Paul R Selvin. Kinesin and dynein move a peroxisome in vivo: a tug-of-war or coordinated movement? Science, 308(5727):1469–1472, June 2005.

[6] Meredith H Wilson and Erika L F Holzbaur. Opposing microtubule motors drive robust nuclear dynamics in developing muscle cells. J. Cell Sci., 125(Pt 17):4158–4169, September 2012.

[7] Yoshimitsu Kanai, Naoshi Dohmae, and Nobutaka Hirokawa. Kinesin transports RNA: isolation and characterization of an RNA-transporting granule. Neuron, 43(4):513–525, August 2004.

[8] Lyudmila Dimitrova-Paternoga, Pravin Kumar Ankush Jagtap, Anna Cyrklaff, Vaishali, Karine Lapouge, Peter Sehr, Kathryn Perez, Simone Heber, Christian Löw, Janosch Hennig, and Anne Ephrussi. Molecular basis of mRNA transport by a kinesin-1-atypical tropomyosin complex. Genes Dev., June 2021.

[9] Yusuke Fukuda, Maria F Pazyra-Murphy, Elizabeth S Silagi, Ozge E Tasdemir-Yilmaz, Yihang Li, Lillian Rose, Zoe C Yeoh, Nicholas E Vangos, Ezekiel A Geffken, Hyuk-Soo Seo, Guillaume Adelmant, Gregory H Bird, Loren D Walensky, Jarrod A Marto, Sirano Dhe-Paganon, and Rosalind A Segal. Binding and transport of SFPQ-RNA granules by KIF5A/KLC1 motors promotes axon survival. J. Cell Biol., 220(1), January 2021.

[10] Alison Emma Twelvetrees, Stefano Pernigo, Anneri Sanger, Pedro Guedes-Dias, Giampietro Schiavo, Roberto A Steiner, Mark P Dodding, and Erika L F Holzbaur. The dynamic localization of cytoplasmic dynein in neurons is driven by kinesin-1. Neuron, 90(5):1000–1015, June 2016.

[11] Atsuko Uchida, Nael H Alami, and Anthony Brown. Tight functional coupling of kinesin-1a and dynein motors in the bidirectional transport of neurofilaments. Mol. Biol. Cell, 20(23):4997–5006, December 2009.

[12] Isabel M Palacios and Daniel St Johnston. Kinesin light chain-independent function of the kinesin heavy chain in cytoplasmic streaming and posterior localisation in the drosophila oocyte. Development, 129(23):5473–5485, December 2002.

[13] Wen Lu, Michael Winding, Margot Lakonishok, Jill Wildonger, and Vladimir I Gelfand. Microtubule-microtubule sliding by kinesin-1 is essential for normal cytoplasmic streaming in drosophila oocytes. Proc. Natl. Acad. Sci. U. S. A., 113(34):E4995– 5004, August 2016.

[14] David D Hackney and Alison E Twelvetrees. The kinesin-1 family: Long-Range transporters. In The Kinesin Superfamily Handbook, pages 15–32. CRC Press, 2020.

[15] Jan O Wirth, Lukas Scheiderer, Tobias Engelhardt, Johann Engelhardt, Jessica Matthias, and Stefan W Hell. MINFLUX dissects the unimpeded walking of kinesin-1. Science, 379(6636):1004–1010, March 2023.

[16] Takahiro Deguchi, Malina K Iwanski, Eva-Maria Schentarra, Christopher Heidebrecht, Lisa Schmidt, Jennifer Heck, Tobias Weihs, Sebastian Schnorrenberg, Philipp Hoess, Sheng Liu, Veronika Chevyreva, Kyung-Min Noh, Lukas C Kapitein, and Jonas Ries. Direct observation of motor protein stepping in living cells using MINFLUX. Science, 379(6636):1010–1015, March 2023.

[17] Johannes F Weijman, Sathish K N Yadav, Katherine J Surridge, Jessica A Cross, Ufuk Borucu, Judith Mantell, Derek N Woolfson, Christiane Schaffitzel, and Mark P Dodding. Molecular architecture of the autoinhibited kinesin-1 lambda particle. Sci Adv, 8(37):eabp9660, September 2022.

[18] Zhenyu Tan, Yang Yue, Felipe da Veiga Leprevost, Sarah E Haynes, Venkatesha Basrur, Alexey I Nesvizhskii, Kristen J Verhey, and Michael A Cianfrocco. Autoinhibited kinesin-1 adopts a hierarchical folding pattern. Elife, 12:2023.01.26.525761, November 2023.

[19] Glenn Carrington, Uzrama Fatima, Ines Caramujo, Tarek Lewis, David Casas-Mao, and Michelle Peckham. A multiscale approach reveals the molecular architecture of the autoinhibited kinesin KIF5A. J. Biol. Chem., (105713):105713, February 2024.

[20] Ahmet Yildiz. Mechanism and regulation of kinesin motors. Nat. Rev. Mol. Cell Biol., pages 1–18, October 2024.

[21] Alison E Twelvetrees, Flavie Lesept, Erika L F Holzbaur, and Josef T Kittler. The adaptor proteins HAP1a and GRIP1 collaborate to activate the kinesin-1 isoform KIF5C. J. Cell Sci., 132(24), December 2019.

[22] John T Canty, Andrew Hensley, Merve Aslan, Amanda Jack, and Ahmet Yildiz. TRAK adaptors regulate the recruitment and activation of dynein and kinesin in mitochondrial transport. Nat. Commun., 14(1):1376, March 2023.

[23] Adam R Fenton, Thomas A Jongens, and Erika L F Holzbaur. Mitochondrial adaptor TRAK2 activates and functionally links opposing kinesin and dynein motors. Nat. Commun., 12(1):4578, July 2021.

[24] David D Hackney, Joelle D Levitt, and Douglas D Wagner. Characterization of *α*2*β*2 and *α*2 forms of kinesin. Biochem. Biophys. Res. Commun., 174(2):810–815, January 1991.

[25] Jacek R Wiśniewski, Marco Y Hein, Jürgen Cox, and Matthias Mann. A “proteomic ruler” for protein copy number and concentration estimation without spike-in standards. Mol. Cell. Proteomics, 13(12):3497–3506, December 2014.

[26] S Hisanaga, H Murofushi, K Okuhara, R Sato, Y Masuda, H Sakai, and Nobutaka Hirokawa. The molecular structure of adrenal medulla kinesin. Cell Motil. Cytoskeleton, 12(4):264–272, January 1989.

[27] David D Hackney, J D Levitt, and J Suhan. Kinesin undergoes a 9 S to 6 S conformational transition. J. Biol. Chem., 267(12):8696–8701, April 1992.

[28] S A Kuznetsov and V I Gelfand. Bovine brain kinesin is a microtubule-activated ATPase. Proc. Natl. Acad. Sci. U. S. A., 83(22):8530–8534, November 1986.

[29] K Dietrich, C Sindelar, P Brewer, K Downing, C Cremo, and S Rice. The kinesin-1 motor protein is regulated by a direct interaction of its head and tail. Proc. Natl. Acad. Sci. U. S. A., 105(26):8938–8943, July 2008.

[30] Hung Yi Kristal Kaan, David D Hackney, and Frank Kozielski. The structure of the kinesin-1 motor-tail complex reveals the mechanism of autoinhibition. Science, 333(6044):883–885, August 2011.

[31] David D Hackney and Maryanne F Stock. Kinesin’s IAK tail domain inhibits initial microtubule-stimulated ADP release. Nat. Cell Biol., 2(5):257–260, May 2000.

[32] J Atherton, M S Chegkazi, E Peirano, L S Pozzer, T Foran, and R A Steiner. Microtubule association induces a mg-free apo-like ADP pre-release conformation in kinesin-1 that is unaffected by its autoinhibitory tail. November 2024.

[33] Kyoko Chiba, Kassandra M Ori-McKenney, Shinsuke Niwa, and Richard J McKenney. Synergistic autoinhibition and activation mechanisms control kinesin-1 motor activity. Cell Rep., 39(9):110900, May 2022.

[34] D L Coy, W O Hancock, M Wagenbach, and J Howard. Kinesin’s tail domain is an inhibitory regulator of the motor domain. Nat. Cell Biol., 1(5):288–292, September 1999.

[35] Ganesh Agam, Christian Gebhardt, Milana Popara, Rebecca Mächtel, Julian Folz, Benjamin Ambrose, Neharika Chamachi, Sang Yoon Chung, Timothy D Craggs, Marijn de Boer, Dina Grohmann, Taekjip Ha, Andreas Hartmann, Jelle Hendrix, Verena Hirschfeld, Christian G Hübner, Thorsten Hugel, Dominik Kammerer, Hyun-Seo Kang, Achillefs N Kapanidis, Georg Krainer, Kevin Kramm, Edward A Lemke, Eitan Lerner, Emmanuel Margeat, Kirsten Martens, Jens Michaelis, Jaba Mitra, Gabriel G Moya Muñoz, Robert B Quast, Nicole C Robb, Michael Sattler, Michael Schlierf, Jonathan Schneider, Tim Schröder, Anna Sefer, Piau Siong Tan, Johann Thurn, Philip Tinnefeld, John van Noort, Shimon Weiss, Nicolas Wendler, Niels Zijlstra, Anders Barth, Claus A M Seidel, Don C Lamb, and Thorben Cordes. Reliability and accuracy of single-molecule FRET studies for characterization of structural dynamics and distances in proteins. Nat. Methods, 20(4):523–535, April 2023.

[36] Eitan Lerner, Anders Barth, Jelle Hendrix, Benjamin Ambrose, Victoria Birkedal, Scott C Blanchard, Richard Börner, Hoi Sung Chung, Thorben Cordes, Timothy D Craggs, Ashok A Deniz, Jiajie Diao, Jingyi Fei, Ruben L Gonzalez, Irina V Gopich, Taekjip Ha, Christian A Hanke, Gilad Haran, Nikos S Hatzakis, Sungchul Hohng, Seok-Cheol Hong, Thorsten Hugel, Antonino Ingargiola, Chirlmin Joo, Achillefs N Kapanidis, Harold D Kim, Ted Laurence, Nam Ki Lee, Tae-Hee Lee, Edward A Lemke, Emmanuel Margeat, Jens Michaelis, Xavier Michalet, Sua Myong, Daniel Nettels, Thomas-Otavio Peulen, Evelyn Ploetz, Yair Razvag, Nicole C Robb, Benjamin Schuler, Hamid Soleimaninejad, Chun Tang, Reza Vafabakhsh, Don C Lamb, Claus Am Seidel, and Shimon Weiss. FRET-based dynamic structural biology: Challenges, perspectives and an appeal for open-science practices. Elife, 10, March 2021.

[37] N Hirokawa, K K Pfister, H Yorifuji, M C Wagner, S T Brady, and G S Bloom. Submolecular domains of bovine brain kinesin identified by electron microscopy and monoclonal antibody decoration. Cell, 56(5):867–878, March 1989.

[38] L A Amos. Kinesin from pig brain studied by electron microscopy. J. Cell Sci., 87 (Pt 1):105–111, February 1987.

[39] Dawen Cai, Adam D Hoppe, Joel A Swanson, and Kristen J Verhey. Kinesin-1 structural organization and conformational changes revealed by FRET stoichiometry in live cells. J. Cell Biol., 176(1):51–63, January 2007.

[40] Fedor V Subach and Vladislav V Verkhusha. Chromophore transformations in red fluorescent proteins. Chem. Rev., 112(7):4308–4327, July 2012.

[41] Jonas Wilhelm, Stefanie Kühn, Miroslaw Tarnawski, Guillaume Gotthard, Jana Tünnermann, Timo Tänzer, Julie Karpenko, Nicole Mertes, Lin Xue, Ulrike Uhrig, Jochen Reinstein, Julien Hiblot, and Kai Johnsson. Kinetic and structural characterization of the Self-Labeling protein tags HaloTag7, SNAP-tag, and CLIP-tag. Biochemistry, 60(33):2560–2575, August 2021.

[42] J T Yang, R A Laymon, and L S Goldstein. A three-domain structure of kinesin heavy chain revealed by DNA sequence and microtubule binding analyses. Cell, 56(5):879–889, March 1989.

[43] Mark A Seeger, Yongbo Zhang, and Sarah E Rice. Kinesin tail domains are intrinsically disordered. Proteins: Struct. Funct. Bioinf., 80(10):2437–2446, July 2012.

[44] D S Friedman and Ronald D Vale. Single-molecule analysis of kinesin motility reveals regulation by the cargo-binding tail domain. Nat. Cell Biol., 1(5):293–297, September 1999.

[45] Michael T Kelliher, Yang Yue, Ashley Ng, Daichi Kamiyama, Bo Huang, Kristen J Verhey, and Jill Wildonger. Autoinhibition of kinesin-1 is essential to the dendrite-specific localization of golgi outposts. J. Cell Biol., 217(7):2531–2547, July 2018.

[46] Maryanne F Stock, J Guerrero, B Cobb, C T Eggers, T G Huang, X Li, and David D Hackney. Formation of the compact confomer of kinesin requires a COOH-terminal heavy chain domain and inhibits microtubule-stimulated ATPase activity. J. Biol. Chem., 274(21):14617–14623, May 1999.

[47] Emil G P Stender, Soumik Ray, Rasmus K Norrild, Jacob Aunstrup Larsen, Daniel Petersen, Azad Farzadfard, Céline Galvagnion, Henrik Jensen, and Alexander K Buell. Capillary flow experiments for thermodynamic and kinetic characterization of protein liquid-liquid phase separation. Nat. Commun., 12(1):7289, December 2021.

[48] Desiree M Baron, Adam R Fenton, Sara Saez-Atienzar, Anthony Giampetruzzi, Aparna Sreeram, Shankaracharya, Pamela J Keagle, Victoria R Doocy, Nathan J Smith, Eric W Danielson, Megan Andresano, Mary C McCormack, Jaqueline Garcia, Valérie Bercier, Ludo Van Den Bosch, Jonathan R Brent, Claudia Fallini, Bryan J Traynor, Erika L F Holzbaur, and John E Landers. ALS-associated KIF5A mutations abolish autoinhibition resulting in a toxic gain of function. Cell Rep., 39(1):110598, April 2022.

[49] Devesh C Pant, Janani Parameswaran, Lu Rao, Isabel Loss, Ganesh Chilukuri, Rosanna Parlato, Liang Shi, Jonathan D Glass, Gary J Bassell, Philipp Koch, Rüstem Yilmaz, Jochen H Weishaupt, Arne Gennerich, and Jie Jiang. ALS-linked KIF5A ΔExon27 mutant causes neuronal toxicity through gain-of-function. EMBO Rep., 23(8):e54234, August 2022.

[50] Benjamin Ambrose, James M Baxter, John Cully, Matthew Willmott, Elliot M Steele, Benji C Bateman, Marisa L Martin-Fernandez, Ashley Cadby, Jonathan Shewring, Marleen Aaldering, and Timothy D Craggs. The smfbox is an open-source platform for single-molecule FRET. Nat. Commun., 11(1):5641, November 2020.

[51] Ronald D Vale, T S Reese, and M P Sheetz. Identification of a novel force-generating protein, kinesin, involved in microtubule-based motility. Cell, 42(1):39–50, August 1985.

[52] S T Brady. A novel brain ATPase with properties expected for the fast axonal transport motor. Nature, 317(6032):73–75, 1985.

[53] Andrew F Macaskill, Johanne E Rinholm, Alison Emma Twelvetrees, I Lorena Arancibia-Carcamo, James Muir, Asa Fransson, Pontus Aspenstrom, David Attwell, and Josef T Kittler. Miro1 is a calcium sensor for glutamate receptor-dependent localization of mitochondria at synapses. Neuron, 61(4):541–555, February 2009.

[54] Kristen J Verhey, D L Lizotte, T Abramson, L Barenboim, Bruce Schnapp, and T A Rapoport. Light chain-dependent regulation of kinesin’s interaction with microtubules. J. Cell Biol., 143(4):1053–1066, November 1998.

[55] F Navone, J Niclas, N Hom-Booher, L Sparks, H D Bernstein, G McCaffrey, and R D Vale. Cloning and expression of a human kinesin heavy chain gene: interaction of the COOH-terminal domain with cytoplasmic microtubules in transfected CV-1 cells. J. Cell Biol., 117(6):1263–1275, June 1992.

[56] Mark A Seeger and Sarah E Rice. Microtubule-associated protein-like binding of the kinesin-1 tail to microtubules. J. Biol. Chem., 285(11):8155–8162, March 2010.

[57] Sumio Terada, Masataka Kinjo, Makoto Aihara, Yosuke Takei, and Nobutaka Hirokawa. Kinesin-1/Hsc70-dependent mechanism of slow axonal transport and its relation to fast axonal transport. EMBO J., 29(4):843–854, February 2010.

[58] R G Elluru, G S Bloom, and Scott T Brady. Fast axonal transport of kinesin in the rat visual system: functionality of kinesin heavy chain isoforms. Mol. Biol. Cell, 6(1):21–40, January 1995.

[59] Sandra Maday, Alison Emma Twelvetrees, Armen J Moughamian, and Erika L F Holzbaur. Axonal transport: Cargo-Specific mechanisms of motility and regulation. Neuron, 84(2):292–309, October 2014.

[60] G S Waldo, B M Standish, J Berendzen, and T C Terwilliger. Rapid protein-folding assay using green fluorescent protein. Nat. Biotechnol., 17(7):691–695, July 1999.

[61] B Ellis, P Haaland, F Hahne, N Le Meur, N Gopalakrishnan, J Spidlen, M Jiang, and G Finak. flowcore: flowcore: Basic structures for flow cytometry data_. R package, 2024.

[62] Antonino Ingargiola, Eitan Lerner, Sangyoon Chung, Shimon Weiss, and Xavier Michalet. FRETBursts: An open source toolkit for analysis of Freely-Diffusing Single-Molecule FRET. PLoS One, 11(8):e0160716, August 2016.

[63] Johannes Schindelin, Ignacio Arganda-Carreras, Erwin Frise, Verena Kaynig, Mark Longair, Tobias Pietzsch, Stephan Preibisch, Curtis Rueden, Stephan Saalfeld, Benjamin Schmid, Jean-Yves Tinevez, Daniel James White, Volker Hartenstein, Kevin Eliceiri, Pavel Tomancak, and Albert Cardona. Fiji: an open-source platform for biological-image analysis. Nat. Methods, 9(7):676–682, July 2012.

[64] Dmitry Ershov, Minh-Son Phan, Joanna W Pylvänäinen, Stéphane U Rigaud, Laure Le Blanc, Arthur Charles-Orszag, James R W Conway, Romain F Laine, Nathan H Roy, Daria Bonazzi, Guillaume Duménil, Guillaume Jacquemet, and Jean-Yves Tinevez. TrackMate 7: integrating state-of-the-art segmentation algorithms into tracking pipelines. Nat. Methods, June 2022.

[65] Richard Evans, Michael O’Neill, Alexander Pritzel, Natasha Antropova, Andrew Senior, Tim Green, Augustin Žídek, Russ Bates, Sam Blackwell, Jason Yim, Olaf Ronneberger, Sebastian Bodenstein, Michal Zielinski, Alex Bridgland, Anna Potapenko, Andrew Cowie, Kathryn Tunyasuvunakool, Rishub Jain, Ellen Clancy, Pushmeet Kohli, John Jumper, and Demis Hassabis. Protein complex prediction with AlphaFold-Multimer. October 2021.

[66] John Jumper, Richard Evans, Alexander Pritzel, Tim Green, Michael Figurnov, Olaf Ronneberger, Kathryn Tunyasuvunakool, Russ Bates, Augustin Žídek, Anna Potapenko, Alex Bridgland, Clemens Meyer, Simon A A Kohl, Andrew J Ballard, Andrew Cowie, Bernardino Romera-Paredes, Stanislav Nikolov, Rishub Jain, Jonas Adler, Trevor Back, Stig Petersen, David Reiman, Ellen Clancy, Michal Zielinski, Martin Steinegger, Michalina Pacholska, Tamas Berghammer, Sebastian Bodenstein, David Silver, Oriol Vinyals, Andrew W Senior, Koray Kavukcuoglu, Pushmeet Kohli, and Demis Hassabis. Highly accurate protein structure prediction with AlphaFold. Nature, 596(7873):583–589, August 2021.

[67] Google colab. https://colab.research.google.com/github/deepmind/alphafold/blob/main/notebooks/AlphaFold.ipynb. Accessed: 2024-12-17.

