## Supplemental Figures for "Kinesin-1 is highly flexible and adopts an open conformation in the absence of cargo"

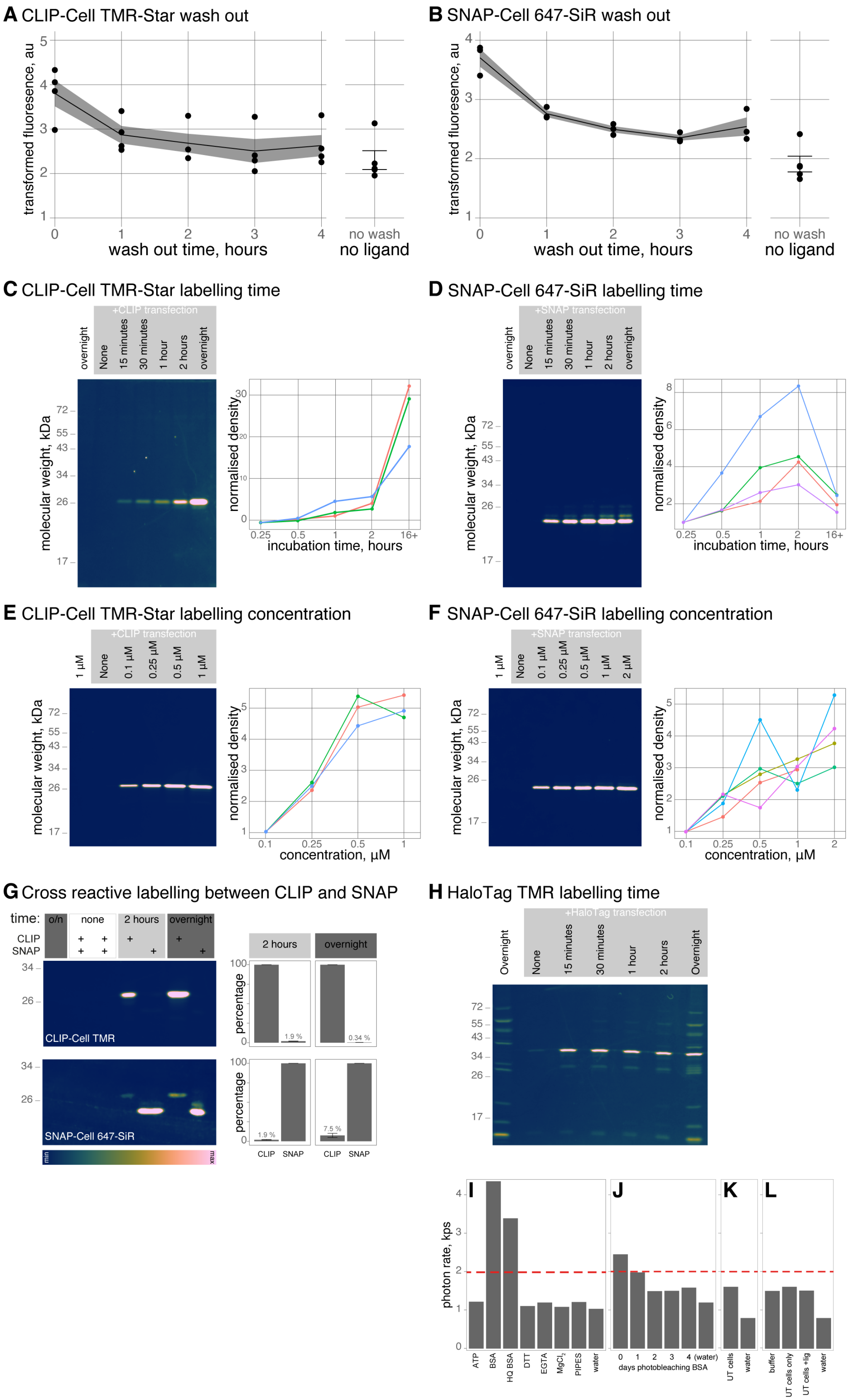


**Figure S1: Optimising labelling of CLIP and SNAP in HEK 293 cells.**
(Continued on following page.)

**Figure S1:**
(A, B) Flow cytometry results showing time taken to wash out unbound CLIP-Cell TMR-Star (A) and SNAP-Cell 647-SiR (B) from non-transfected HEK 293 cells. Fluorescence signal from untransfected and unlabelled cells is shown for comparison. At least three biological replicates were acquired for each time point.
(C, D) In-gel fluorescence of cell lysate from HEK 293 cells transfected with CLIP or SNAP alone and incubated with CLIP-Cell TMR-Star (C) or SNAP-Cell 647-SiR (D) for the indicated time periods. Quantified signal intensity for biological replicates is shown, with labelling intensity normalised to the first time point. At least three biological replicates were acquired for each time point. The batlow scientific colour map was applied to all fluorescent gel images to indicate intensity, and is illustrated in panel G.
(E, F) In-gel fluorescence of cell lysate from HEK 293 cells transfected with CLIP or SNAP alone and incubated with CLIP-Cell TMR-Star (E) or SNAP-Cell 647-SiR (F) at the indicated concentrations for 2 hours or overnight for SNAP and CLIP respectively. Quantified signal intensity for biological replicates is shown, with labelling intensity normalised to the first concentration point. At least three biological replicates were acquired for each concentration. The batlow scientific colour map was applied to all fluorescent gel images to indicate intensity, and is illustrated in panel G.
(G) Cross reactivity of CLIP and SNAP ligands for their counter tag was measured by in-gel fluorescence of cell lysate from HEK 293 cells, transfected with either CLIP-tag or SNAP-tag and incubated with ligands for time periods as indicated. Even overnight labelling with CLIP-Cell TMR-Star produced minimal cross reactive labelling with the SNAP-tag (0.34%), however SNAP-Cell 647-SiR did have more significant cross reactivity with CLIP-tags over extended time periods (7.5%). Consequently CLIP-Cell TMR-Star was incubated on transfected HEK cells overnight, whilst SNAP-Cell 647-SiR was limited to 2 hours before washing and lysate collection.
(H) In-gel fluorescence revealed significant non-specific labelling of the HaloTag TMR ligand when incubated with HEK 293 cells for longer than 30 minutes, both in HaloTag transfected (grey box) and untransfected cells. The batlow scientific colour map was applied to all fluorescent gel images to indicate intensity, and is illustrated in panel G.
(I-L) Background photon rates (kilo photons per second, kps) observed in components of the smFRET assay buffer. Red dotted line marks the threshold for the photon rate, above which background photons begin to interfere with burst analysis. When each buffer component was assayed individually (I), BSA (even high quality BSA) was found to be a significant source of background photon bursts when illuminated by ALEX. Photobleaching the BSA stock over several days (J) in a lab-made broad wavelength illumination box, reduced this background to acceptable levels. Cell lysate from untransfected cells diluted 10,000 fold (comparable to typical assay conditions in smFRET) does not add considerably to the background when compared to water alone (K). In the optimised final smFRET assay conditions (L) the kinesin smFRET assay buffer (made with photo bleached BSA, buffer), untransfected HEK 293 cell lysate diluted in assay buffer (cells only), and diluted lysate from cells treated with both fluorophores and washed out according to the optimised labelling conditions (cells +lig), all displayed background photon rates low enough for reliable smFRET analysis. Data in L was collected routinely to ensure sample and buffer integrity prior to data collection.

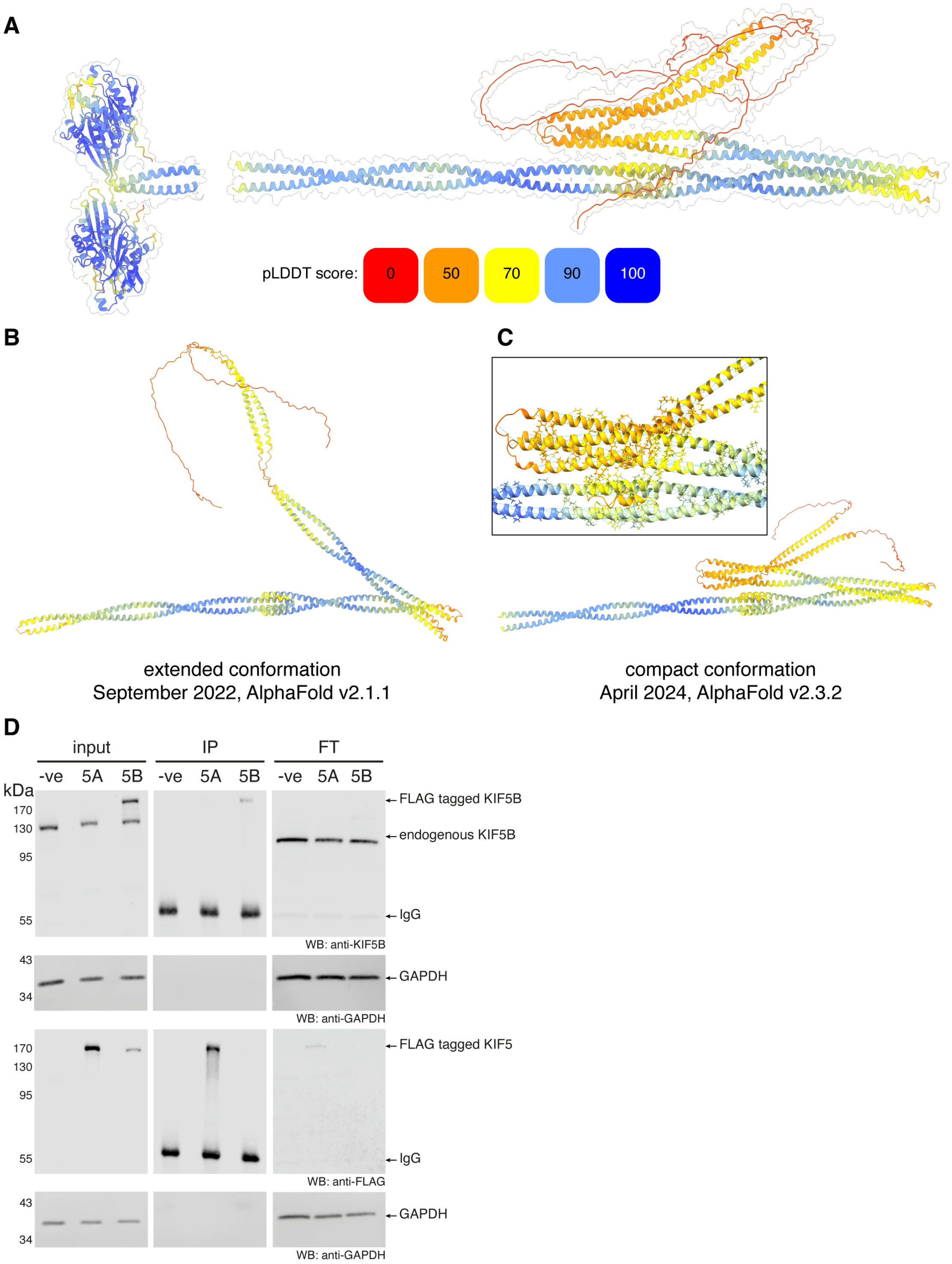


**Figure S2: AlphaFold predictions of the kinesin-1 stalk have compacted over release versions.**
(A) The same AlphaFold2 model shown in Figure 2B, coloured by pLDDT score.
(B, C) AlphaFold structures of human KIF5C(413-957), encompassing the stalk and tail, generated in 2022 (B) and 2024 (C), coloured by pLDDT score. Inset in panel C highlights the increased hydrogen bonding and electrostatic interactions driving compaction of coiled coils across CC1-CC4.
(D) Exogenous KIF5A does not co-immunoprecipitate endogenous KIF5B from HEK 293 cells. Our CLIP-SNAP expression vector (pCLAP) also contains a C-terminal FLAG tag. HEK 293 cells were transfected with mouse CLIP-KIF5A-SNAP-FLAG (5A) or human CLIP-KIF5B-SNAP-FLAG (5B). The overexpressed proteins were immunoprecipitated with anti-FLAG antibody. Samples were separated by SDS-PAGE and probed by western blot (WB) using the antibodies: rabbit anti-KIF5B, mouse anti-GAPDH and rabbit anti-FLAG. Although KIF5A was readily immunoprecipitated, no corresponding endogenous KIF5B band was observed. Tagged KIF5B did not overexpress to the same level as KIF5A, but with the endogenous KIF5B antibody a similar result could be observed. Samples: input, cleared cell lysate; IP, immunoprecipitation; FT, flow through. Blots representative of three biological replicates.

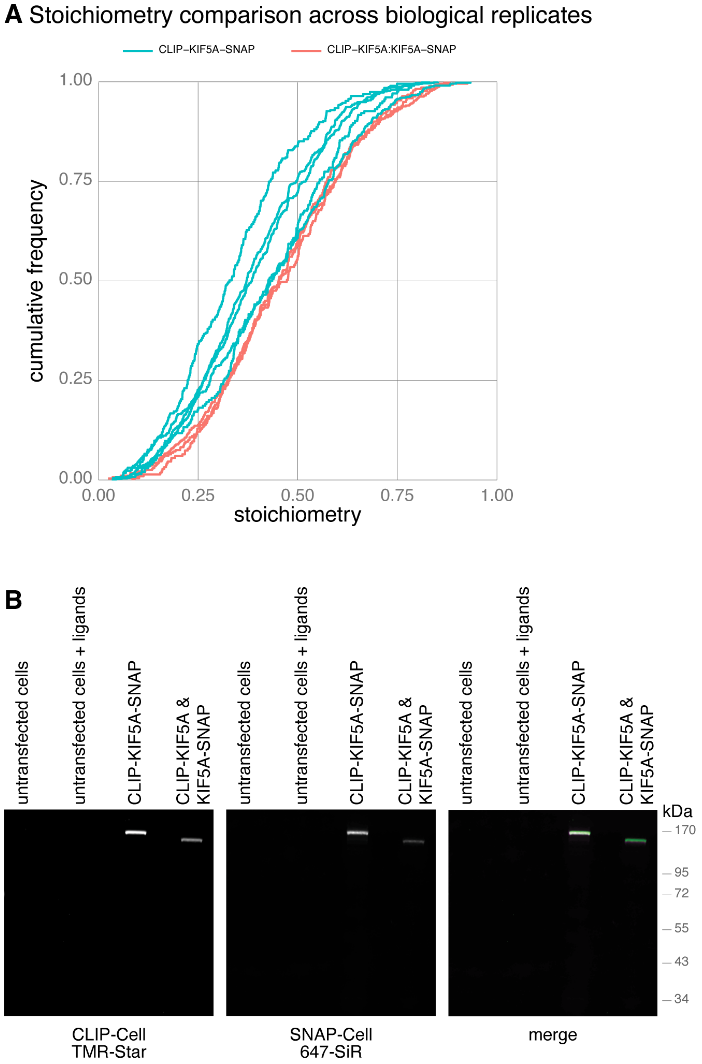


**Figure S3: Variable stoichiometry across biological replicates for homodimers of CLIP-KIF5A-SNAP.**
(A) Cumulative frequency plots of stoichiometry for individual biological replicates of CLIP-KIF5A-SNAP homodimers (blue) compared to CLIP-KIF5A & KIF5A-SNAP heterodimers (red). Homodimers show more variability in the stoichiometry of labelling, due to the variable labelling efficiency of CLIP compared to SNAP. The low proportion of observed heterodimer molecules poses challenges for collecting data in this regime, despite a more consistent stoichiometry from eliminating labelling variability.
(B) In-gel fluorescence of cell lysate from HEK 293 cells transfected with CLIP-KIF5A-SNAP or CLIP-KIF5A & KIF5A-SNAP heterodimers, and incubated with CLIP-Cell TMR-Star and SNAP-Cell 647-SiR. The merged fluorescence panel (right) shows 647-SiR in magenta and TMR-Star in green. CLIP-KIF5A or KIF5A-SNAP are a lower molecular weight as they are shorter by the equivalent of one tag (~20 kDa) compared to CLIP-KIF5A-SNAP.

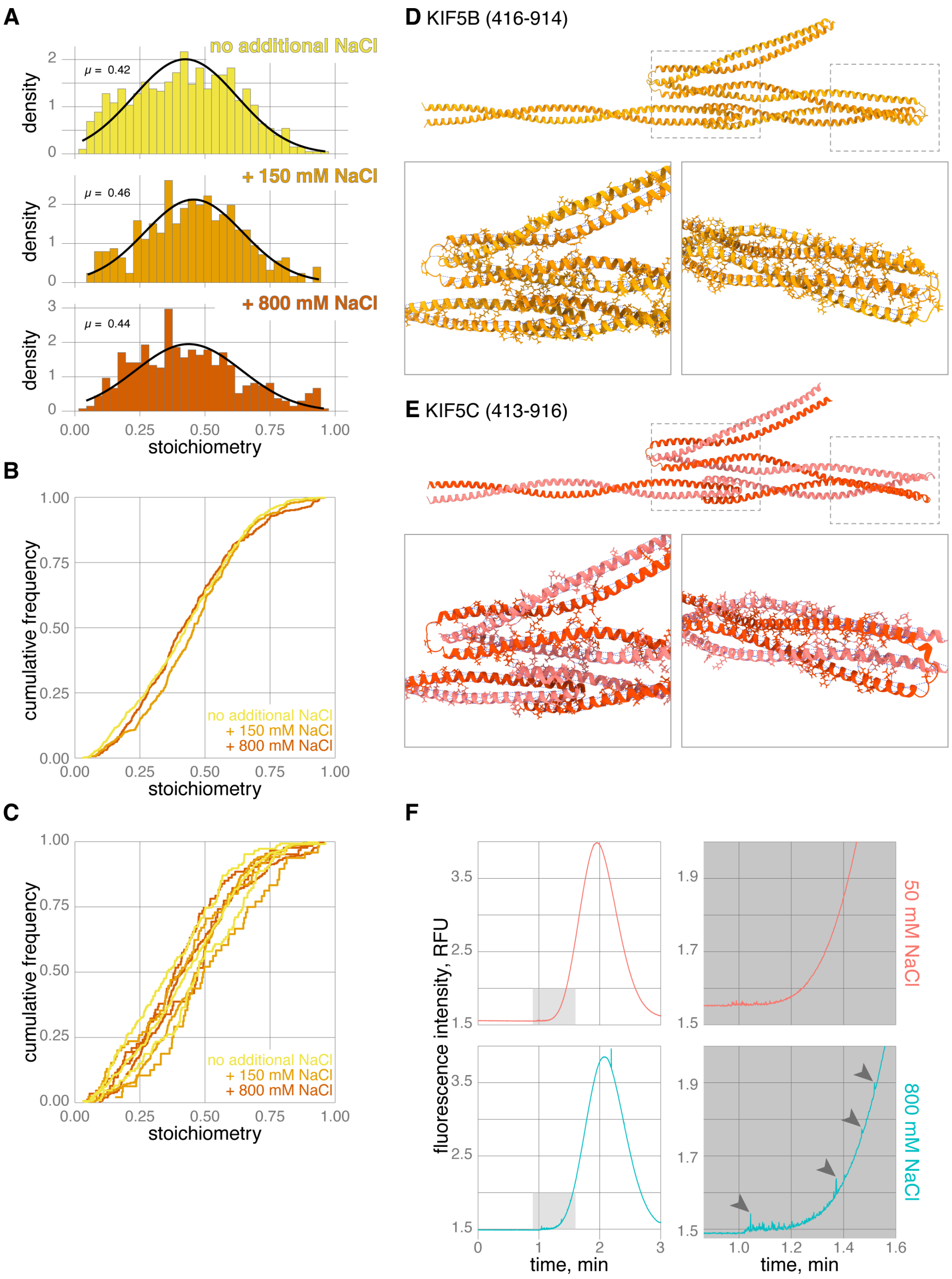


**Figure S4: Stoichiometry is insensitive to NaCl concentration.**
(A - C) Histograms (A), cumulative frequency plots of pooled data (B) and biological replicates (C), comparing the stoichiometry of CLIP-KIF5A-SNAP samples in 0 (yellow), 150 (orange) or 800 mM (brown) additional NaCl.
(D, E) AlphaFold2 predictions of the stalk domains of KIF5B (D) and KIF5C (E) with expanded regions below highlighting hydrogen bonds and electrostatic interactions involved in stacking the coiled coils domains against each other in Hinge 1 (right) and across CC1-4 incorporating Hinge 2 (left).
(F) Representative Taylorgrams for CLIP-KIF5A-SNAP in 50 mM (red) or 800 mM (blue) additional NaCl. Grey box is expanded to the right to show spikes of aggregated protein highlighted by arrowheads.

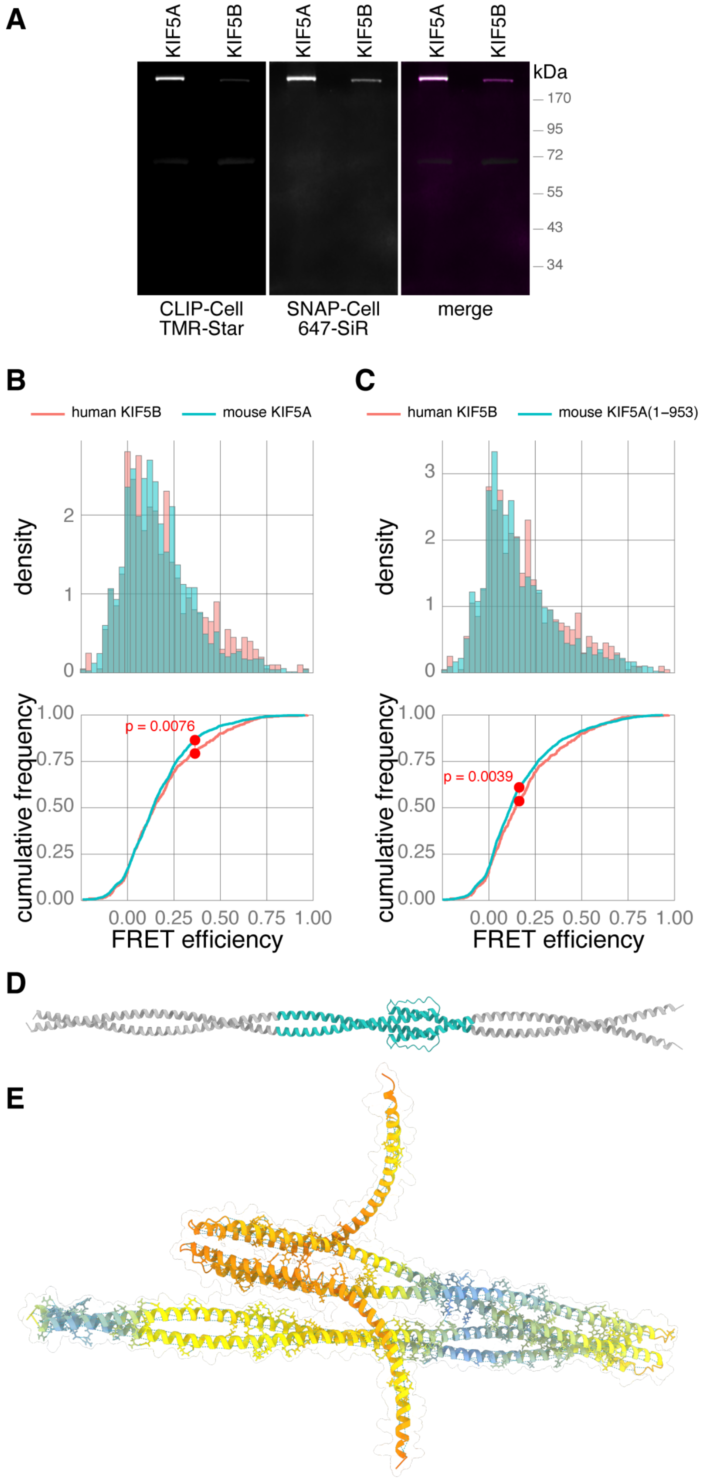


**Figure S5: Consistency in FRET efficiency across Kinesin-1 isoforms.**
(A) In-gel fluorescence of cell lysate from HEK 293 cells transfected with CLIP-KIF5A-SNAP or CLIP-KIF5B-SNAP, and incubated with CLIP-Cell TMR-Star and SNAP-Cell 647-SiR. The merged fluorescence panel (right) shows 647-SiR in magenta and TMR-Star in green. KIF5B consistently had lower expression than KIF5A in this expression system, which may reflect tighter regulation of protein levels for an isoform expressed endogenously in HEK 293 cells.
(B, C) Histograms and cumulative frequency plots comparing the FRET efficiency of human KIF5B (red) to full length mouse KIF5A (B) or KIF5A(1-953) (C), truncated to be equivalent in length to KIF5B. Full length KIF5A has a slightly higher FRET efficiency compared to KIF5B, which would be anticipated due to its longer length, whereas KIF5A(1-953) is almost identical.
(D) An AlphaFold2 prediction of CC1-CC2 of KIF5A, with deleted residues 505-610 (spanning the retrograde loop), highlighted in turquoise.
(E) An AlphaFold2 prediction of KIF5A tail (411-1027) with residues 505-610 deleted (residues 921-1027 are hidden for clarity). The structure is coloured by pLDDT score. The predicted structure of the deletion mutant shows that the coiled-coil structure of CC1-CC2 of the stalk is retained with deletion, but the removal of the retrograde loop removes interactions between CC2 to CC3, with knock-on effects for the positioning of CC4.

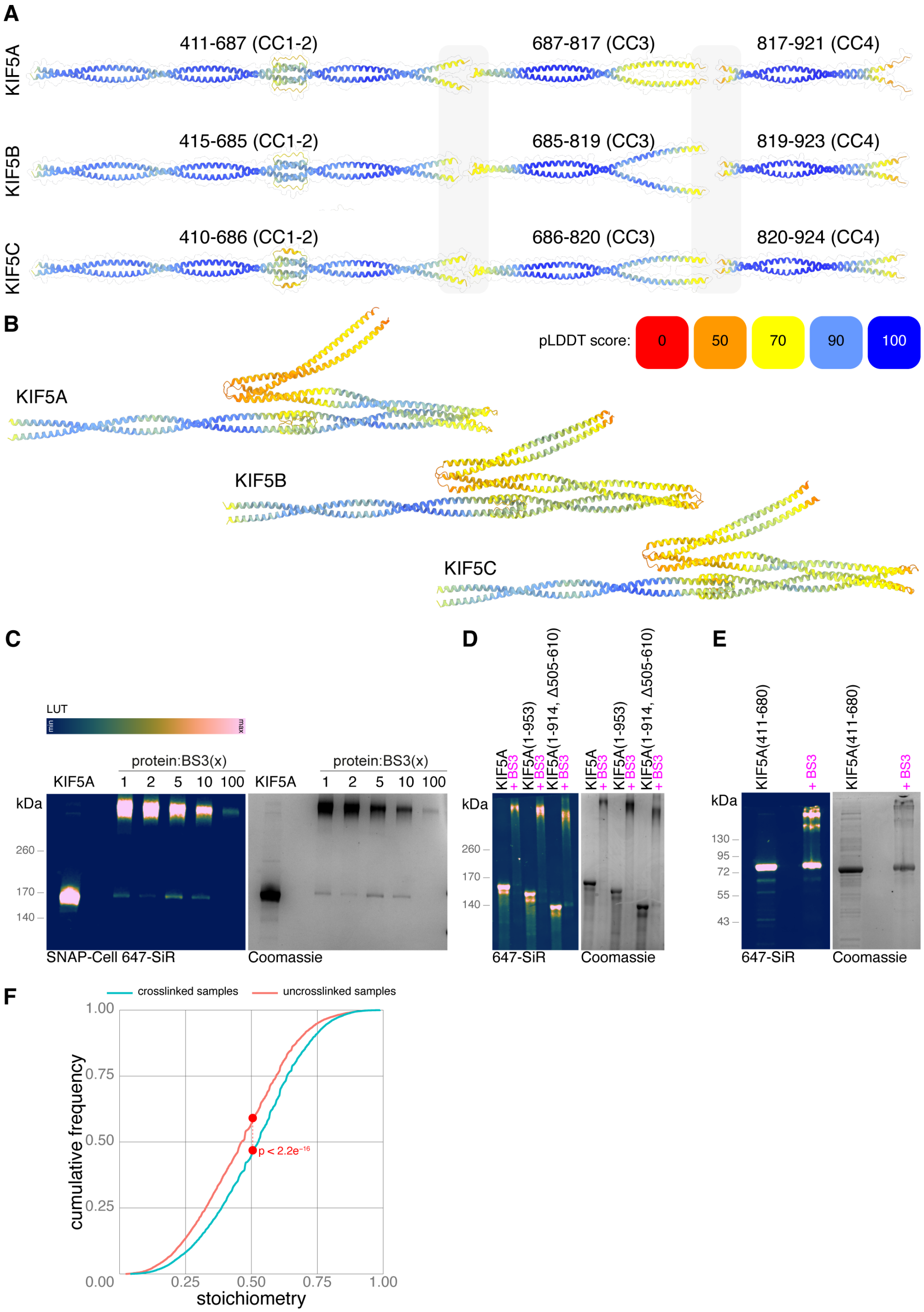


**Figure S6: AlphaFold predictions of domains suggest conformational flexibility.**

(Continued on following page.)

**Figure S6:**

(A, B) AlphaFold2 predictions of kinesin-1 stalk domains either individually (A) or as a complete stalk (B). Boundaries between domains for A were defined by predicted aligned errors (PAE) domains, and centred on the unstructured regions in full tail domain predictions (namely the linker, hinge 1 and hinge 2 in Figure 2B). This creates three coiled coil regions from left to right: CC1-CC2 (packed together by the retrograde loop), CC3 and CC4. Grey boxes highlight phase breakdown of coiled-coils. Predictions illustrated in B were modelled as the complete tail domain, including the C-terminal intrinsically disordered region, but this region was omitted from the figure for clarity.
(C) In-gel fluorescence of 647-SiR labelled CLIP-KIF5A-SNAP (left) and Coomassie stain of the same (right) purified from HEK 293 cells and treated with various ratios (by mass) of the crosslinking reagent BS3. The amount of BS3 is indicated above the lane and compared to the uncrosslinked sample labelled 'KIF5A'. A ratio of 1:1 was used for onward experiments, as little uncrosslinked KIF5A remained and higher amounts of BS3 trapped kinesin in the well, resulting in lower band intensities at high molecular weight. The batlow scientific colour map was applied to the fluorescent gel image to indicate intensity.
(D, E) Example protein samples from Figure 6 separated by SDS-PAGE and imaged by in-gel fluorescence of 647-SiR labelled SNAP or by Coomassie stain. Full length KIF5A and mutants (as indicated, D) or a CC1-CC2 KIF5A fragment (E) purified from HEK 293 cells and optionally treated with a 1:1 ratio by mass of the crosslinking reagent BS3. The batlow scientific colour map was applied to the fluorescent gel images to indicate intensity, and is illustrated in panel C.
(F) A cumulative frequency plot comparing the stoichiometry of crosslinked (blue) and uncrosslinked (red) samples. Stoichiometry always shifted to the right in crosslinked samples, strongly implying that BS3 had a tendency to quench fluorescence, particularly of the SNAP-tag.
